# From average transient transporter currents to microscopic mechanism – A Bayesian analysis

**DOI:** 10.1101/2023.10.31.565026

**Authors:** August George, Daniel M. Zuckerman

**Affiliations:** Department of Biomedical Engineering, Oregon Health and Science University, Portland, OR, USA

## Abstract

Electrophysiology studies of secondary active transporters have revealed quantitative, mechanistic insights over many decades of research. However, the emergence of new experimental and analysis approaches calls for investigation of the capabilities and limitations of the newer methods. We examine the ability of solid-supported membrane electrophysiology (SSME) to characterize discrete-state kinetic models with *>* 10 rate constants. We use a Bayesian framework applied to synthetic data for three tasks: to quantify and check (i) the precision of parameter estimates under different assumptions, (ii) the ability of computation to guide selection of experimental conditions, and (iii) the ability of SSME data to distinguish among mechanisms. When the general mechanism – event order – is known in advance, we show that a subset of kinetic parameters can be “practically identified” within *∼*1 order of magnitude, based on SSME current traces that visually appear to exhibit simple exponential behavior. This remains true even when accounting for systematic measurement bias and realistic uncertainties in experimental inputs (concentrations) are incorporated into the analysis. When experimental conditions are optimized or different experiments are combined, the number of practically identifiable parameters can be increased substantially. Some parameters remain intrinsically difficult to estimate through SSME data alone, suggesting additional experiments are required to fully characterize parameters. We additionally demonstrate the ability to perform model selection and determine the order of events when that is not known in advance, comparing Bayesian and maximum-likelihood approaches. Finally, our studies elucidate good practices for the increasingly popular, but subtly challenging, Bayesian calculations for structural and systems biology.

## Introduction

Transporters are a type of biological molecular machine that help regulate cellular homeostasis by pumping molecules across a membrane and maintaining ion gradients.^1,2^ As such, transporters play an essential role in cellular processes, such as the uptake of nutrients and expelling of waste. These protein systems operate in a stochastic molecular environment which suggests they could exhibit some degree of stochasticity in their mechanism, a concept recently emerging as ‘pathway heterogeneity’,^3–6^ but which has been implicit in reports of non-integer stoichiometry of secondary active transport over many years.^7–11^ In a recent example, analysis of the small *Escherichia coli* multidrug transporter^12^ points to utilization of different sequences of biochemical steps that enable a wide range of behaviors including 2:1 and 1:1 transport stoichiometry.^4,13^ Despite significant study, the precise mechanisms of transport – i.e., the states visited, the allowed order(s) of transitions, and the associated rate constants – remain difficult to characterize.

Traditional electrophysiology techniques^14,15^ have performed an essential role in many important discoveries related to membrane transport, such as determining the kinetics of ion channels^16,17^ and neuronal transmission,^15^ and have a long-standing history within the transporter research community.^18,19^ They provide dynamic information regarding the transport of ions across a membrane under various external conditions, which can be used to estimate specific kinetic parameters such as net transport rates. Solid-supported membrane electrophysiology (SSME),^20,21^ extends traditional electrophysiology methods by introducing a fixed membrane embedded with reconstituted proteoliposomes that are perturbed under different external concentrations. This approach yields an averaged and aggregated transient current trace with a large signal relative to the noise, and is stable across multiple perturbation concentrations in sequence, such as the three-stage reversal assay.^22^ However, it is unknown how well SSME can recapture microscopic information from the generated macroscopic dataset. That is, how much information is contained in these datasets?

We address the question of inferring mechanistic details from SSME data using Bayesian inference (BI). We are not aware of prior applications of BI to transporter parameter or mechanism inference. BI provides a well-established and robust framework for estimating model parameters and their uncertainties from noisy datasets, as well as for distinguishing among models. Briefly, Bayesian inference^23^ is a powerful statistical method that generates a “posterior” probability *distribution* of model parameters given the dataset and prior beliefs about the model. The approach is computationally expensive compared to alternative frequentist methods such as maximum likelihood estimation (MLE),^23^ but provides a comprehensive posterior distribution that contains most likely parameter estimates, uncertainties, and correlations, in addition to enabling comparison among models. In practice, the posterior distribution of parameters is estimated using Markov chain Monte Carlo (MCMC),^24–26^ which may prove challenging to converge with many systems of interest that are multidimensional and embody complex sampling landscapes, akin to rough energy landscapes. Here we utilize a recently developed sampling approach that combines powerful ideas from physics (annealing) and from machine-learning (normalizing flows)^27^ to overcome convergence challenges.

Building on related work for ion channels^28,29^ and methods developed for computational systems biology,^30,31^ and molecular biophysics,^32–34^ we implement a Bayesian inference data analysis pipeline to address the gap in knowledge about membrane transporters mechanisms and solid-supported membrane electrophysiology datasets. To our knowledge, this is the first application of Bayesian inference for transporter mechanism. We use ordinary differential equations (ODE) models of transporter reaction cycles and SSME-like conditions to generate synthetic data. The synthetic data enables a cost-effective and controlled environment with known ground truth values for methods development, but at the cost of potential simplifications and biases not found in experimental data. Synthetic data also can be used hand-in-hand with experimentation to design more informative experiments, as detailed below.

In this study, we demonstrate the effectiveness of BI for three important tasks in analyzing SSME data: parameter and uncertainty estimation; optimization of experimental conditions; and inference of mechanism, i.e., model selection. Using a (known) default model and data set, after validating our pipeline, we examine the practical identifiability issue: the precision with which each of *>*10 model rate constants can be determined. We consider the important, but often overlooked, effects of uncertainties in “known” experimental conditions, namely, species concentrations. We next consider an array of experimental conditions to ascertain which experiments and experiment combinations are the most informative for parameter inference. Finally, we use a Bayesian model selection strategy for four 1:1 transport cycles to ascertain the data required to distinguish the underlying model when that is not known in advance. We also examine MLE results for model selection.

We find that the Bayesian inference pipeline is well suited for these key tasks in SSME-based transporter research, generating highly informative posterior distributions across the various models and synthetic assays studied. The results reveal that most model rate constants can be determined within *∼*1 order-of-magnitude precision with suitable data – starting from a six-order-of-magnitude range – but that certain rate-constants are intrinsically less identifiable and may require independent measurements. The experiment optimization ranking reveals which SSME conditions contain the most information – typically with those with large pH/concentration changes, as well as combinations of different assays. Also, we found that high-information datasets enable distinguishing among four possible mechanistic models, whereas low-information datasets could not. These results illustrate the likely utility of BI in studying transporter mechanisms, giving insight into optimal experimental design, and suggesting experiments that may be able to fully determine the reaction pathways of membrane transporters. Comparison of MLE calculations for model selection with BI results shows that certain MLE algorithms, properly tuned, can be successful but that BI may be a more robust approach overall. Finally, we also describe our experiences – and lessons learned – in MCMC sampling for BI calculations.

## Results

We present results for transporter parameter inference from SSME data, for comparison of the informativeness of different experimental protocols for parameter inference, and for determination of the transport mechanism (event order). We focus on 1:1 exchangers (antiporters) based on the classical alternating access paradigm,^35,36^ described via a system of ordinary differential equations (ODEs) along with Gaussian noise. Throughout, we use synthetic data for 1:1 ion:substrate exchange generated using the ODE model for “Cycle 1” (Figure 2) described in Methods. The model has 12 rate constants, of which 11 are independent due to the thermodynamic cycle constraint.^37^ Synthetic data enables validation of the calculations, and by itself represents a significant challenge for MCMC sampling. Alternative mechanisms for 1:1 transport, i.e., different event orders, are considered below.

**Figure 1:**
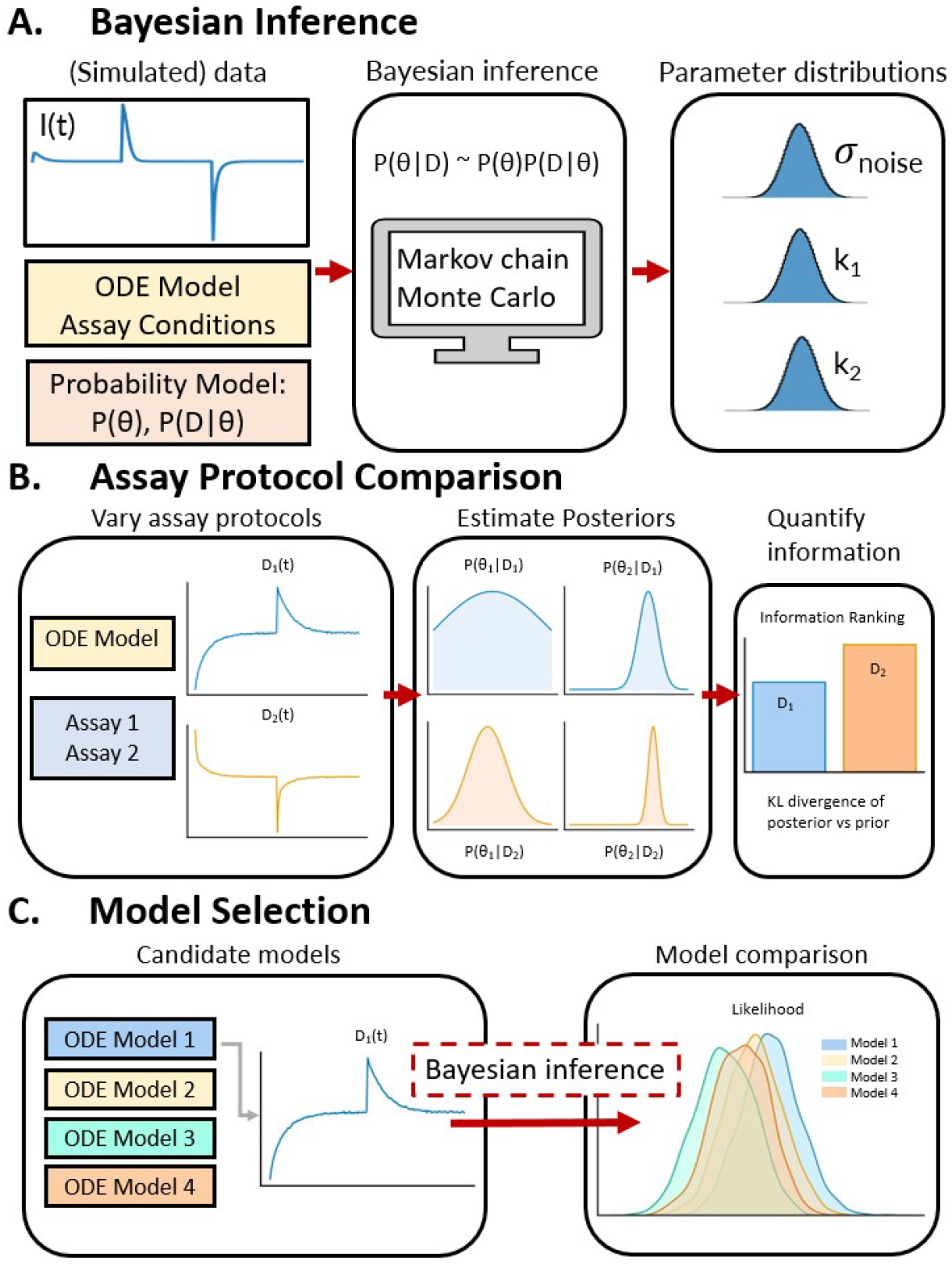
Overview of data analysis pipelines for transporter research. **(A)** Bayesian inference is used with synthetic SSME assay data to generate parameter estimate distributions for reaction rate constants and other nuisance parameters (e.g., noise variance). **(B)** Different assay conditions are simulated from the same model, with the KL divergence between the posterior and priors used to approximate the information contained in each dataset. **(C)** The likelihoods of different transporter reaction cycles are compared using the same dataset for model selection.

**Figure 2:**
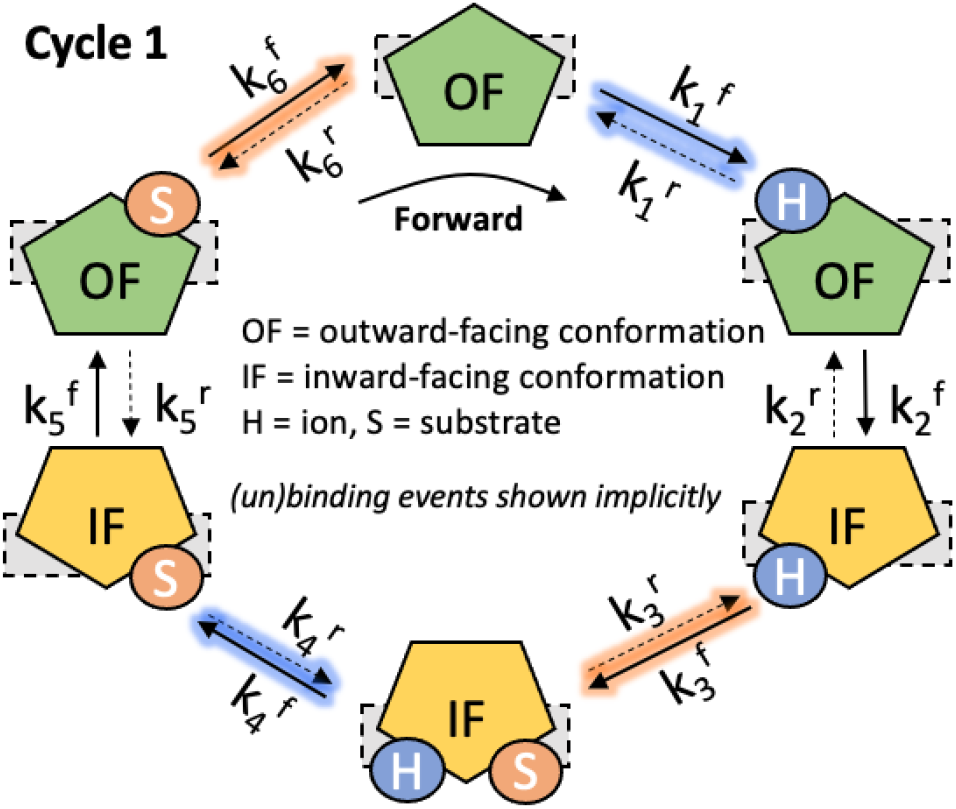
A 1:1 transport cycle (antiport/exchange). Starting from the top-most fully unbound outward-facing (OF) state and proceeding clockwise, an outside proton (H) binds, followed by conformational eversion to the inward-facing (IF) state, then binding of an inside substrate (S), and so on. Note that binding and unbinding events are implicit in the state transitions. The time evolution of each state is described by a system of ODEs given in Methods, with rate constants shown at transitions. This “Cycle 1” model is used to generate the synthetic data used in the study.

The SSME current data models a three-stage protocol,^22^ shown in Figure 3, with current values from the activation and reversal phases used in analysis. Different experimental protocols considered here correspond to activation by different external substrate concentration and pH values.

**Figure 3:**
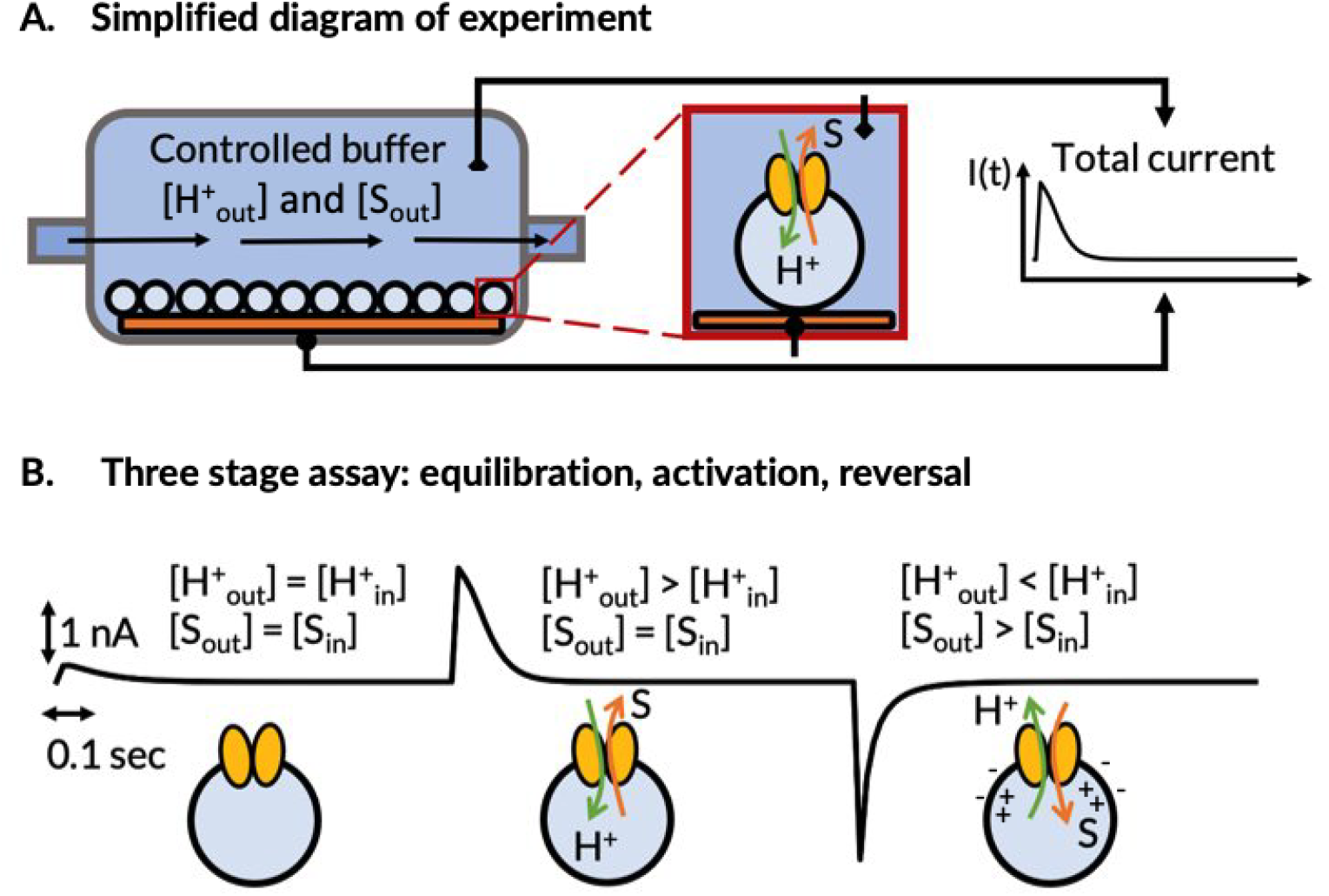
Schematic of SSME experiments. In solid-supported membrane electrophysiology (SSME) experiments generally (A), transporters embedded on liposomes transfer ions under a driving gradient, inducing a current. The bottom panel (B) shows a three-stage reversal assay: After an initial equilibration phase with matching concentrations inside and outside liposomes, the concentrations of the external species are perturbed in an activation phase to induce a transient current of ions across the membrane, and the external perturbation is subsequently reversed.

Importantly, our BI calculations start from extremely non-informative prior ranges: rate constants are assumed to be unknown – and equally likely – within a range of six orders of magnitude. This breadth of parameter space makes the calculations challenging, but is important in avoiding implicit foreknowledge of the true parameter values. MCMC sampling is performed withe the pocoMC package, which uses an unbiased annealing process in combination with the “normalizing flows” approach.^27^

### Parameter estimation, comparing limited and more realistic experimental uncertainties

An central issue in the analysis of experimental data is accounting for measurement noise and potential bias, not only for the output of the experiment – e.g., the current in electrophysiology – but also in “known” control parameters such as the concentrations of molecular and ionic species. Previous work analyzing calorimetry data has demonstrated the importance of accounting for possible measurement error in concentrations and, in some cases, shown that Bayesian inference evidently can correct concentrations when a physical model for the system is known.^33,38^

We therefore characterized the parameters for the Cycle 1 antiporter (Figure 2 using both a simplified and more realistic set of parameters. We first considered a 12-parameter set (12D model) consisting of the 11 independent rate constants of Cycle 1 plus a single “nuisance” parameter *σ* for the noise in the current measurement. We also studied a 16D model which included four additional nuisance parameters: *f*_bias_ representing a systematic linear bias in the current to account for uncertainty intrinsic to SSME measurements,^39^ and *f*_protein_,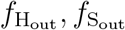 representing uncertainties in concentrations of transporters, ion (or proton, H), and substrate (S) following our prior work.^33^ The four additional parameters are represented as dimensionless scaling factors; see Methods.

Our initial analysis is based on the “Experiment 1” dataset, which represents a SSME three-stage protocol run a single time. See Methods for conditions chosen. As will be seen, the Experiment 1 dataset is relatively uninformative by comparison to other sets.

The estimated posterior distribution shown in Fig. 4, projected as one-dimensional marginals. Overall, only a few parameters can be determined precisely from the simple Experiment 1 dataset analyzed; however, below, we will see significant improvement in parameter identifiability with improved datasets. When BI is able to find distributions which are narrow compared to the prior (full ranges shown), those include the known true parameter values. Furthermore, general consistency among independent MCMC replicas suggests the posteriors are well sampled.

**Figure 4:**
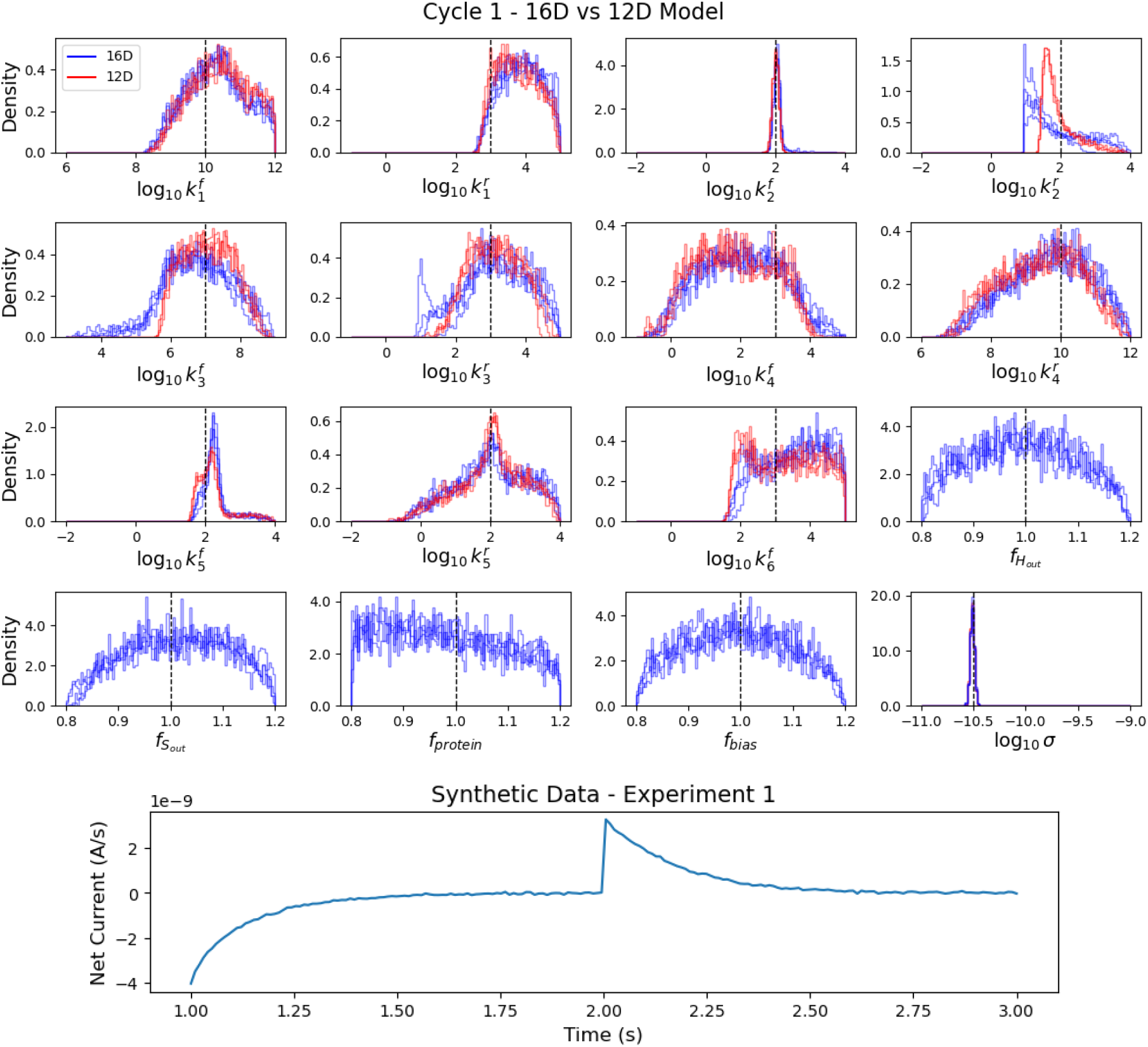
Parameter inference, comparing the effect of simple and more realistic parameterizations (Experiment 1). The estimated parameter distributions for two different model assumptions across replicas are shown, with ground truth values (vertical lines) for reference. Here the 12D model (red) parameters consists of 11 rate constants and Gaussian noise standard deviation. The 16D model (blue) additionally includes uncertainty in the initial assay concentrations and an measurement bias term. The horizontal range for each distribution matches the width of the uniform prior, notably six orders of magnitude for rate constants. Only data from “Experiment 1” were analyzed here, as shown in the bottom panel for reference. The 12D model used 3 sampling replicas, while the 16D model used 4 sampling replicas.

Most important here is the comparison between 12D and 16D parameterizations, which is indicative of the ‘cost’ of including realistic nuisance parameters for experimental uncertainties, as well as the feasibility of MCMC sampling in the more more complex case. Although distributions for the rate constants are slightly broader in the 16D case, we see that inclusion of the additional nuisance parameters does not dramatically degrade parameter identifiability. Further, the sampling is slightly worse in the 16D case, but the replicas are consistent enough overall to distinguish which parameters are practically identifiable, i.e., determined within *∼*1 order of magnitude. These results are further supported by examining the sum of standard deviations, which indicates a modest decrease the estimated precision of the model parameters when using the 16D model (shown in Figure SI 13).

Going forward, we employ only the more realistic 16D parameterization.

### Information quantification across experimental conditions

Next we examine optimizing the design of experiments by quantifying the gain in information and parameter precision when different datasets are employed, reflecting different experimental conditions and/or replicates. This is done by using the 16D model described above to generate datasets from several different assay protocols (detailed in SI) based on four different sets of experimental conditions. As before, multiple replicas of a sequential Monte Carlo^27^ Bayesian inference algorithm with broad uniform priors and a standard Gaussian-noise log-likelihood are run for each dataset.

Posterior distributions comparing different datasets are shown in Fig. 5, and provide strong evidence for improved parameter identifiability compared with the dataset considered previously in Figure 4. As data more from more diverse experimental conditions is included, the posterior marginals become narrower. In the best case examined, the combination of Experiments 1+2+3+4 leads to the effective identification (within *∼*1 order of magnitude) of six out of 11 rate constants. Fig. 5 also shows the effect of including ‘technical replicates’ of Experiment 1, which leads to better precision for many rate constants, but only to a slight degree. Generally, rate constants for conformational transitions were better determined than (un)binding rates, and there was not a major difference between ion and substrate parameter identifiability. It is notable that two-dimensional posterior marginals (Figure SI 12) show strong correlations among the IF binding and unbinding steps, effectively identifying the dissociation constants for steps 3 and 4. In the present data, by comparing MCMC replicates for the same experimental protocol, we see that sampling is not fully converged, but it does appear to be sufficient to support conclusions about parameter identifiability.

**Figure 5:**
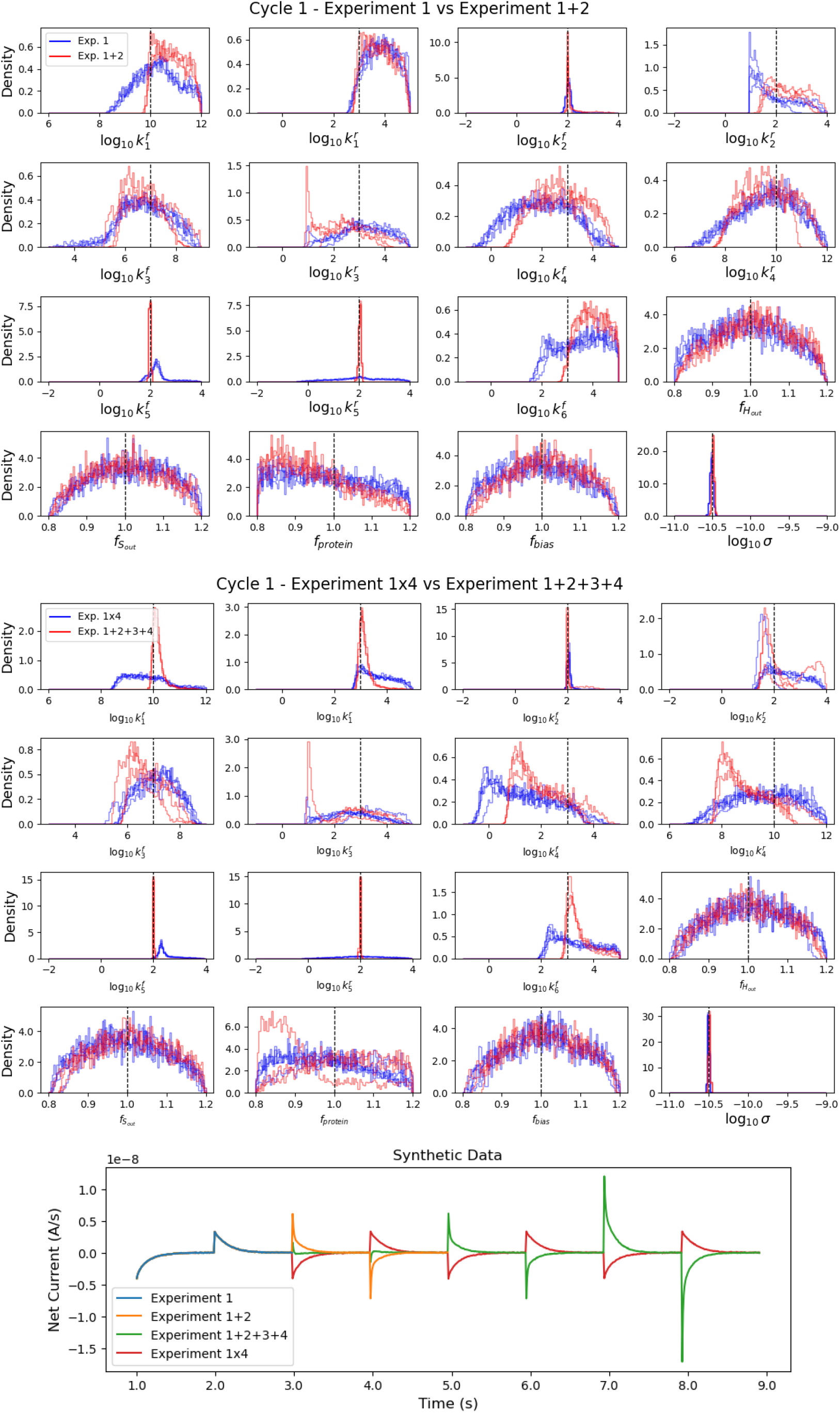
Marginal Posterior Distribution Comparisons: The estimated parameter distributions for varying different synthetic assay protocols across replicas are shown, with ground truth values for reference. **(Top)** A single three-stage perturbation assay (blue, Exp. 1) is compared against a sequence of two three-stage perturbations assays (red, Exp. 1+2). The introduction of a second unique three-stage perturbation assay into the dataset yields a significant reduction in estimate variance. The experiment 1 protocol used 4 sampling replicas, while the experiment 1+2 protocol used 3 sampling replicas. **(Middle)** A single three-stage perturbation assay (blue, Exp. 1×4) is replicated four times in sequence, and compared against a sequence of four unique three-stage perturbations assays (red, Exp. 1+2+3+4). The introduction of additional three-stage perturbation assays into the dataset yields a significant reduction in estimate variance as compared to technical replicas. The experiment 1×4 protocol used 5 sampling replicas, while the experiment 1+2+3+4 protocol used 3 sampling replicas. **(Bottom)** Traces of the synthetic SSME-like data used. We note that for the combined experiments 1-4 dataset, the perturbation corresponding the experiment 2 is third sequentially (see SI).

We can quantify information content of the various protocols via the KL divergence of the posterior distribution relative to the prior distribution^40^ for each dataset. This effectively estimates the information gained from our dataset as compared to an uninformative prior. Further, we calculate the sum of standard deviations of each parameter in the marginalized posterior distributions for each dataset to quantify the precision of the parameter estimates. These results are shown in Fig. 6. As expected from the posterior distributions, the combination of unique experimental conditions contain the more information and higher precision than a single experiment alone or technical replicas. Also, these results show a correlation between the ‘information gained’ and practical identifiability of parameters.

**Figure 6:**
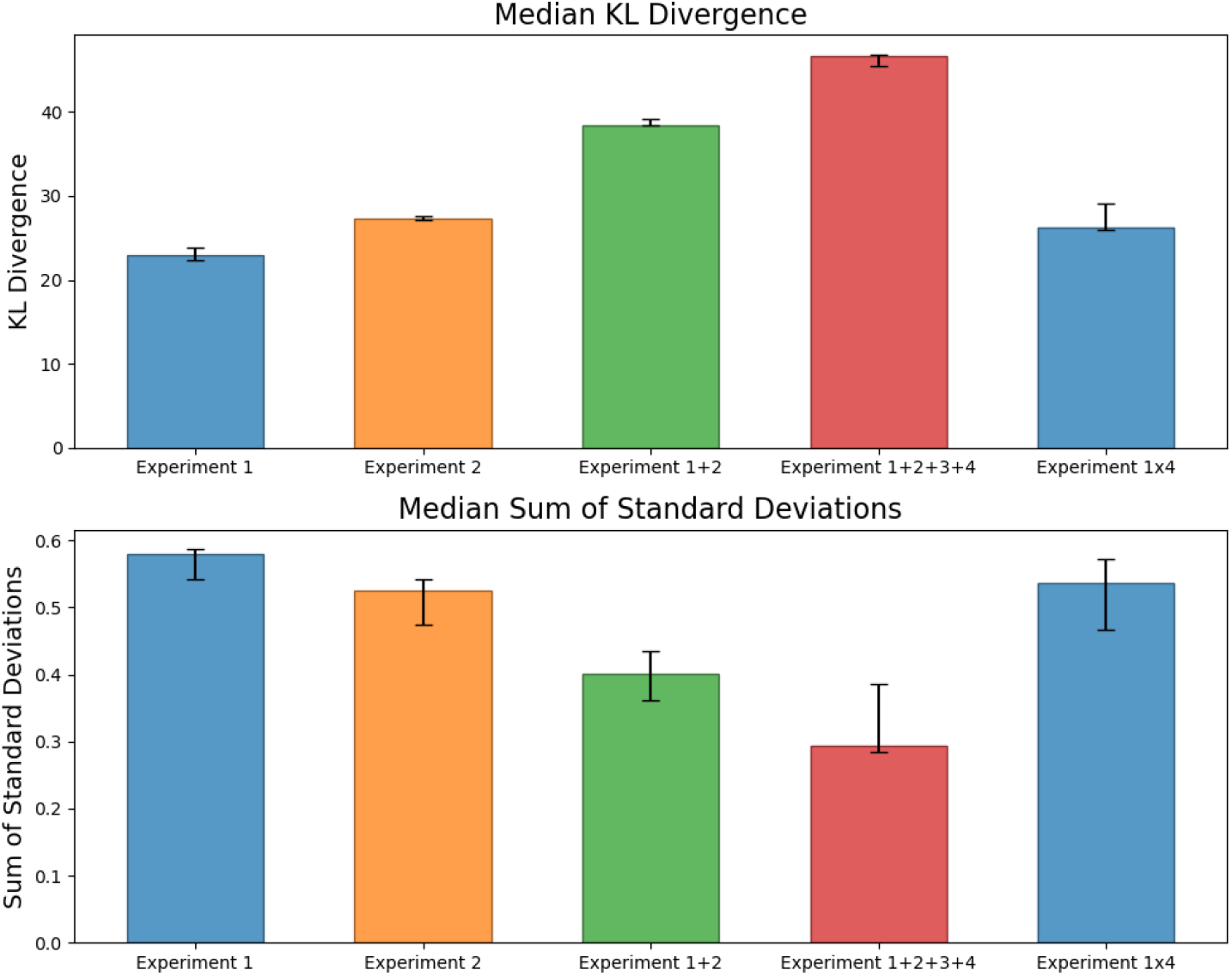
Quantifying information across experimental protocols. **(Top)** The median KL divergence of the posterior and uninformative uniform priors across multiple replicas, with error bars spanning the minimum to maximum values. **(Bottom)** The median sum of standard deviations of the parameter distributions, across multiple replicas are shown with error bars denoting the range from minimum to maximum. The introduction of additional experiments significantly increases the information gained as compared to a single experiment or technical replicas. The sum of standard deviations is correlated with the KL divergence and indicates an improvement in parameter precision as additional experiments are introduced. We note that KL divergence is dimensionless, but can be expressed in terms of nats. Similarly, the sum of standard deviations does not correspond to a physical quantity, but rather is a composite dimensionless measure used to compare the relative estimated precision across varying conditions.

### Transporter mechanism identification (model selection)

We now examine whether SSME current traces are sufficient to select among different event orders, i.e., mechanisms. We consider the four 1:1 antiport pathways shown in Figure 7, which includes the previously considered cycle (Fig. 2) along with three additional mechanisms; see Methods for details. For comparison to BI calculations, we use maximum likelihood estimation (MLE)^23^ with the differential evolution algorithm^41^ which was found empirically to perform well on a test (simpler) dataset after hyper-parameter tuning (see SI). We examine the log-likelihoods for each model and data set, using both Bayesian and MLE methods as shown in Fig 8. Finally, these results are compared with the estimated marginal likelihood (i.e. model evidence) values generated during sampling.

**Figure 7:**
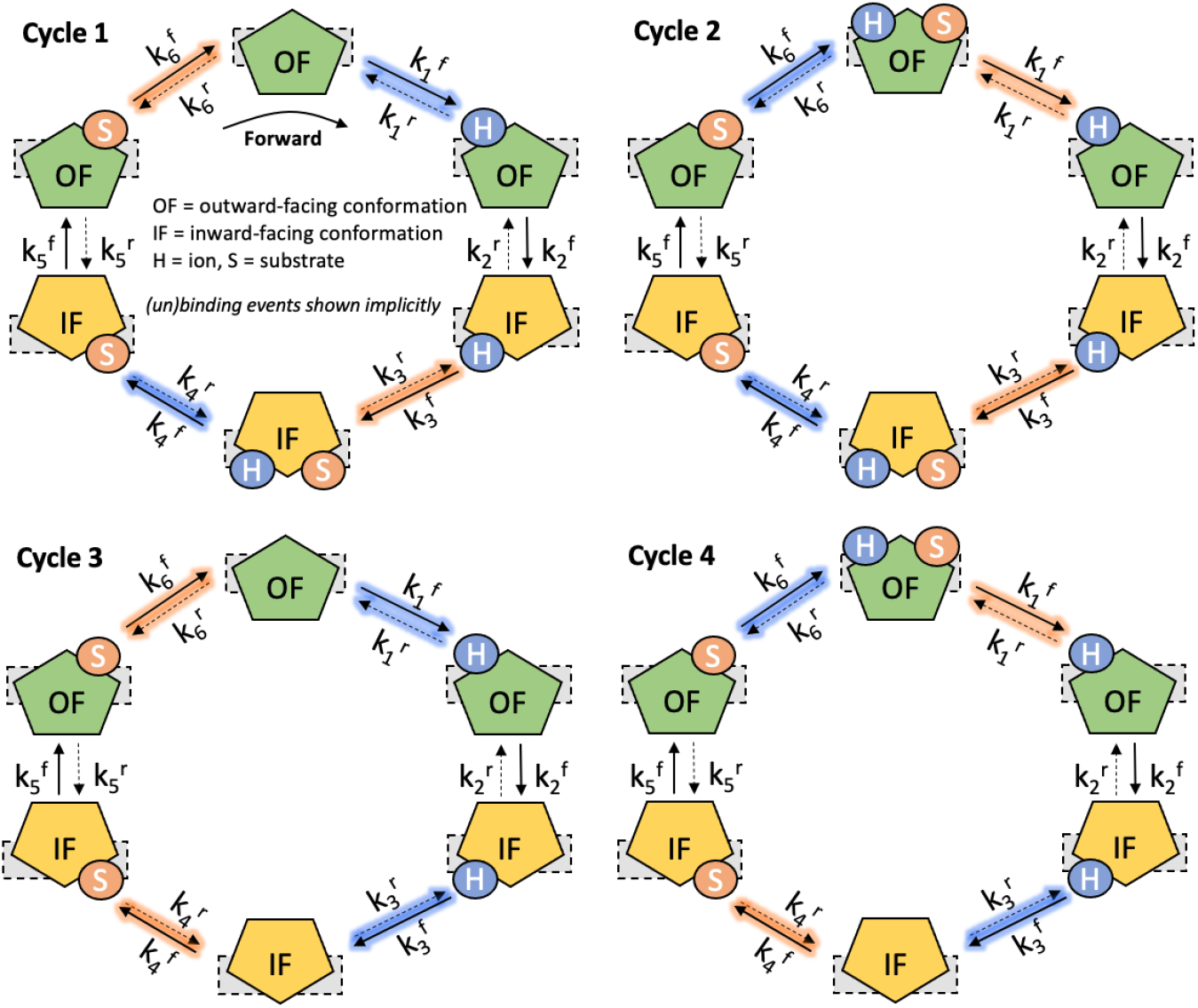
Four Tightly Coupled 1:1 Antiporter Reaction Cycles. Each cycle transports an ion (H) and a substrate (S) in opposite directions via alternating access (outward-facing, OF, and inward-facing, IF) conformations. Each network has unique set of six reaction states which represent different reaction event orders. The ion (un)binding reactions are highlighted in blue, with the substrate (un)binding reactions are highlighted in orange, with each cycle having a unique combination of ion and substrate reactions. The binding and unbinding (i.e. dissociation) events are shown implicitly for improved visual clarity.

**Figure 8:**
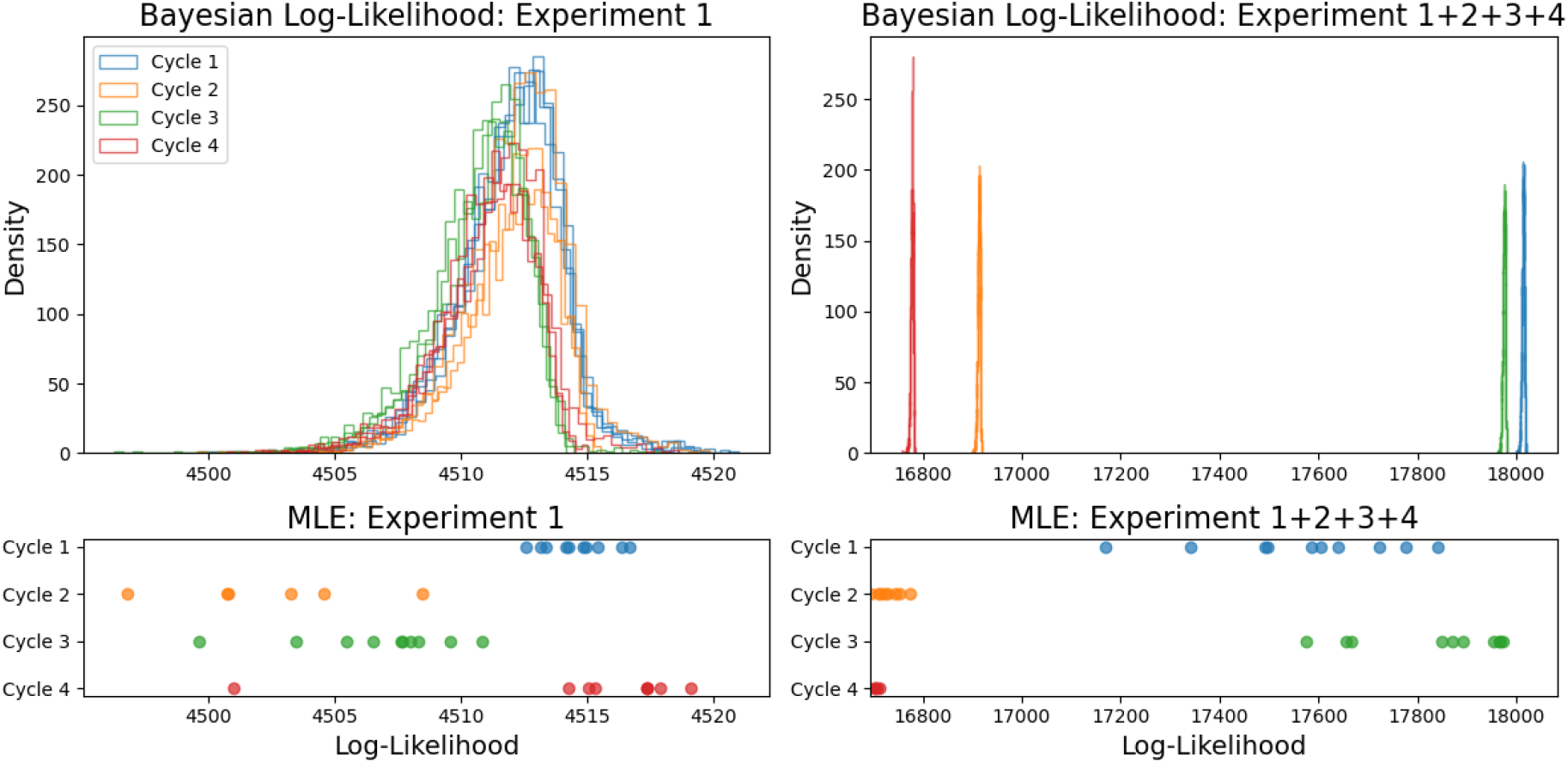
Transporter Mechanism Selection (Top Row) The Bayesian log-likelihood distributions of four transporter reaction cycles using less (left) and more (right) informative datasets, across multiple replicas. The less informative dataset yields a large overlap between the model likelihood distributions, while the more informative dataset yields a large divergence between the likelihoods - suggesting a single most likely model given the data. **(Bottom Row)** The maximum log-likelihood distributions of four transporter reaction cycles using less (left) and more (right) informative datasets, across multiple replicas. This approach generates a range of likelihood values across models and datasets, but fails to reliably converge on the maximum likelihood. We note that the ‘Cycle 1’ (blue) transporter mechanism is used as a ground truth and is expected to have the largest likelihood.

Inferring which mechanism generated a given dataset, the task of “model selection”, turns out to require the most informative dataset explored for parameter inference. The Bayesian inference results show a general overlap between each reaction cycle log-likelihood distribution when the less informative dataset (Expt 1) is used, but a clear separation between distributions when the informative dataset (Expt 1+2+3+4) is used. The results correctly identify ‘Cycle 1’ as the most likely model by many orders of magnitude when using the informative dataset, but with the uninformative dataset all the models have a similarly high likelihood. The maximum likelihood results fail to reliably converge to the expected maximum likelihood across the conditions studied, despite tuning of hyperparameters as described in the SI. Additional Bayesian posterior distributions for the cycles and datasets are shown in Figure SI 5-11.

Our results comparing the log-likelihoods are further supported when examining the marginal likelihood (i.e. model evidence) for each model under an informative and uninformative data set, as shown in SI Figure 14. We find that the marginal likelihoods for each model have small relative differences when using an uninformative dataset. In contrast, when an informative data set is used, there is a significant relative difference between the marginal likelihoods for each model – with the ground truth model having the highest evidence.

## Discussion

Understanding detailed mechanisms of molecular processes is a central goal of modern structural biology, and here we have applied Bayesian inference (BI) to study the mechanism of driven biological transport, apparently for the first time. We have examined the recently developed solid-supported membrane electrophysiology (SSME) approach, which is designed for (relatively) high throughput study of transporters with with high signal-to-noise.^22,39^ Synthetic SSME data allowed us study one of the simplest realistic systems, a 1:1 antiporter based on ground-truth reference values, and also enabled facile assessment of the potential value of additional experiments; as detailed below, BI computations even for synthetic data represent a significant challenge. Our approach accounted for important, often overlooked facets of experimental uncertainty, including both concentration measurement uncertainty^33^ and potential systematic measurement bias for electrophysiological currents.^39^

Our study revealed both striking findings and challenges for quantifying mechanism in transporters. First, it is surprising how much information about individual model parameters resides in SSME data, despite the simple-exponential-like visual appearance of the data. The BI results (Figure 5) show that six out of 11 rate constants of a 1:1 transport model are “identifiable” – estimated within an order of magnitude (or less) – starting from a highly permissive six-order-of-magnitude prior range for every rate constant. Further, the correlation structure of the BI posterior (Figure SI 12) for the remaining unidentified rate constants implicitly determines two dissociation constants – i.e., rate-constant ratios – and provides effective guidance on which parameters should be determined from independent experiments. With the experimental data used, the conformational transition rate constants generally are better determined than on- and off-rates, but neither substrate nor ion (un)binding rates have a clear advantage for identifiability.

In comparing mechanistic pathways in a “model selection” task, the results (Figure 8) suggest that a well-chosen set of experiments can enable successful model selection – more than a single experiment or technical replicates – revealing the sequence of mechanistic steps. These results demonstrate the potential of Bayesian inference to determine an unknown transporter mechanism from SSME data, or to eliminate unlikely candidate mechanisms. While we modeled each cycle individually, the Bayesian framework is compatible and could be extended for mixed and hierarchical models (i.e., combinations of multiple pathways). As such, this study provides a foundation to systematically determine the mechanisms of more complex transporter cycles such as the proposed model for EmrE.^13^

The technical challenges involved with both BI and maximum likelihood (MLE) computations for synthetic datasets were significant. The less-than-ideal agreement among replica MCMC runs (Figure 5) highlights the sampling challenge despite our use of a sophisticated and highly parallelized MC sampling approach (pocoMC)^27,42^ which required approximately 12 hours of computing per replicate. During the course of this study, we examined quite a few MCMC methods, none of which could provide the performance of pocoMC; for reference we show a comparison to the affine-invariant ensemble sampler^43^ in Figure SI 20-21. Likewise, we examined a series of maximum likelihood methods, and most of the methods failed to optimize our systems even after tuning of hyper-parameters (Figure SI 15-18); the MLE data shown in Figure 8 is from the differential evolution algorithm^41^ which was the best performer. We found that the Bayesian inference methods were the most computationally expensive, but had a comparable efficiency (number of likelihood calculations per second) as MLE methods such as differential evolution (see Figure SI 19), while generating full posterior distributions rather than point estimates. On the whole, our data on what might be considered a simple model with synthetic data clearly demonstrate the technical challenges of Bayesian inference^30,44^ and highlight the need for careful evaluation of MCMC sampling and MLE optimization.

While the synthetic SSME data studied here was motivated by experimental assay conditions and parameters^22,45^ and we accounted for concentration uncertainty and bias, experimental data will present new challenges. Higher ion:substate stoichiometry will introduce additional parameters and pathways which may require model simplification.^46^ Issues of transporter polarity and uniformity across liposomes, time delays from the mechanics of fluid mixing, and the effects of membrane capacitive coupling^39^ may require additional “nuisance” parameters in models. Future work will aim to address these challenges through close collaboration with SSME-experimentalists and an extension of our current modeling framework. We emphasize nevertheless that a Bayesian framework “self reports” on parameter and model uncertainty and hence can indicate when addition data is needed for BI and/or whether independent measurements of certain parameters are required. Although the present study was limited to SSME-like data, the generic approach should be applicable to many other measurements.

## Conclusion

This work has attempted to shed new light on the study of transporter proteins through the use of Bayesian inference (BI) and information theory to compare mechanisms, determine rate constants for elementary steps, and guide experiment design. We performed a systematic BI study for 1:1 ion:substrate secondary active transporters based on the emerging solid-supported membrane electrophysiology (SSME) experimental approach. Because the task of transporter mechanistic inference involves a combination of complex mechanisms, experiments under a potentially wide range of conditions, and numerically demanding analysis, we employed synthetic SSME data to understand experimental information content and data requirements. Encouragingly, we found that a majority of rate constants for individual mechanistic steps could be determined from well-designed SSME experiments and that the order of events could also be inferred. Although we found that BI was more reliable for model selection than maximum-likelihood approaches, overall our results also underscored the challenges of sampling for BI and the need for careful assessment. Taken as a whole, the study is a necessary step towards a more complete understanding of complex transporter reaction mechanisms, enabling effective model comparison, precise parameter estimation, and informed experimental design.

## Methods

In the following section we describe the computational methods used to model membrane transporters, simulate SSME assays, and perform data analysis. A more comprehensive description of the methods with implementation details can be found in the Supplemental Information.

### Modeling Membrane Transporters

Building on the foundational work by Mitchell^35^ and Jardetzky^36^ we model membrane transport using the alternating access model. We represent these models using biochemical networks, with the dynamics determined by ordinary differential equations governed by mass action kinetics.^47^ There are four idealized 1:1 antiporter models^48^ that transport a single ion (H) and substrate (S) in opposite directions across the membrane. With two conformations, outward-facing (OF), and inward-facing (IF), there are six reaction states, with a unique set of reactions and conformational states distinguishing the models as illustrated in Fig. 7. These differences lead to a unique order of reaction events. For example, in cycles 1 and 3, 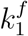 corresponds to an ion binding rate constant, but in cycles 2 and 4, 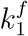 corresponds to a substrate unbinding rate constant. The different physical processes arise from the different states used in the model, such as with an unbound outward-facing state used for cycles 1 and 3, and a doubly bound outward-facing conformation in cycles 2 and 4. We primarily use the ‘Cycle 1’ model in this study, with the exception of the model selection results which utilize all four models. The rate constants use an dynamic Arrhenius-like formulation, with membrane voltage found empirically to have a negligible effect on the simulated data for the conditions studied (see SI Figure 1-2). To ensure detailed balance, one of the twelve rate constants is necessarily constrained by thermodynamic consistency.^37^ Their governing ordinary differential equations along with further simulation and parameter details are described in the SI.

### Generating Synthetic SSME-like Data

As described previously, SSME experiments^20^ consist of many proteoliposomes with reconstituted proteins deposited on a solid membrane. This system is placed in a bath of chemical species and when perturbed creates a gradient that drives ion transport across the membrane that is measured. After an initial equilibration stage, the concentrations are perturbed (i.e., activated), and the system relaxes to a new steady-state condition. In a three-stage reversal assay,^22^ the external concentrations are adjusted back to the initial values in the final stage, switching the gradient and driving transport in the opposite direction (i.e., reversal). Importantly, due to the stability of SSME, multiple assays can be done in sequence under different perturbation amounts. Figure 3 shows an idealized diagram of an SSME experiment, with synthetic data traces shown in Figures 4, 5, and SI Figure 3.

We numerically simulate these assays by integrating the transporter cycle ODEs, with conditions changed for equilibration, activation, and reversal stages at t=0, 1, and 2 seconds (and additional stages as needed). During each assay stage, the external concentrations of the ion and subsrate in the bath solution are held fixed. We use a stiff ODE solver (CVODES)^49^ with a low tolerance to improve numerical stability. The net current is calculated from the change in internal ion concentrations of a single liposome – converting from the change in molar concentration to current, and multiplying by the number of liposomes in the experiment.

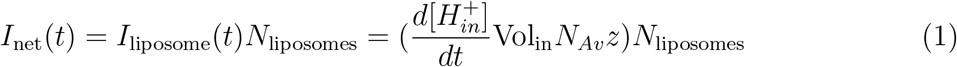

where Vol_in_ is the internal volume of a single liposome embedded with transporters, *N*_*Av*_ is Avogadro’s constant, *z* is the elementary charge of an *H*^+^ ion, and *N*_liposomes_ is the total number of liposomes in the SSME assay. Here we assume a known constant liposome volume and number liposomes, as well as uniformity across the aggregate of liposomes. We note that while our data generating function in Eq. 1 is a relatively simple macroscopic observable with few parameters, the change in the ion concentration 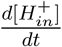 is coupled to the other differential equations that have latent parameters (i.e. microscopic rate constants).

### Bayesian Inference and Parameter Estimation

The predicted SSME-like data is used in the Bayesian pipeline to estimate the probability of the model parameters given the data, as given by Bayes’ theorem:

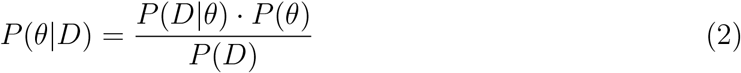

where:

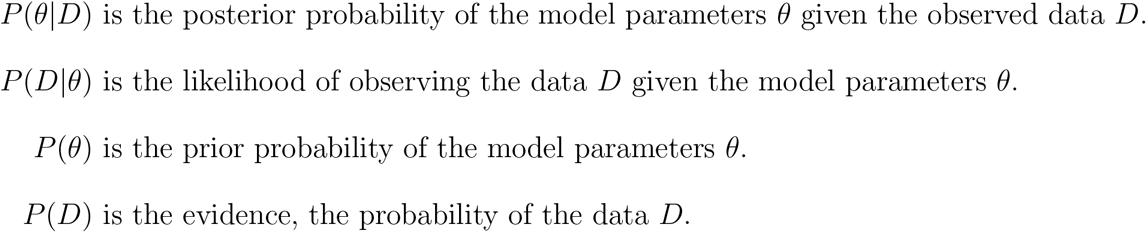

We use a standard log-likelihood function, which assumes Gaussian experimental noise:

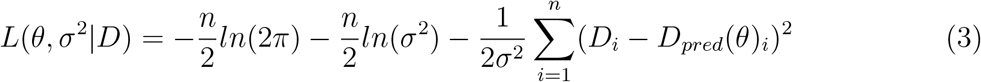

Here our synthetic observation data, *D* = (*D*_1_, *D*_2_, …, *D*_*n*_), is generated from Eq. 1 by solving the governing ODEs with a set of reference (‘ground truth’) reaction rate constants, compartment volumes, number of liposomes, and initial concentrations, and then adding Gaussian white noise. During inference calculations, one set of trial parameters at a time, i.e., the vector *θ*, is used to generate the ‘predicted’ data, *D*_*pred*_, and evaluated in (3) using a candidate Gaussian noise variance *σ*^2^. Additional nuisance parameters, for concentration uncertainties and experimental bias, are subsumed into *θ*.

The reaction rate constants are represented on a log10 scale with wide uniform prior distributions covering six orders of magnitude to reduce bias in our analysis. Further, we incorporate several nuisance parameters for the noise variance, initial species concentrations, and a biasing factor, each with uniform priors.

We use the “pocoMC” package for Monte Carlo sampling, which combines a physically motivated annealing protocol with machine-learning accelerated preconditioned Monte Carlo sampler.^27^ The annealed importance sampling framework,^50,51^ in the context of BI, employs a pseudo-temperature parameter to transition from effectively infinite temperature (uniform sampling) to the posterior distribution of interest. At each intermediate “temperature” a neural network is trained using normalizing flows^52^ via specialized autoencoder neural networks^53,54^ to simplify the geometry of the sampling space. We found this method to have improved performance over alternative Bayesian and maximum likelihood estimation methods (see SI).

### Information Quantification and Experiment Optimization

For experiment optimization and recommendation, we are primarily interested in screening for protocols that yield high information data and reduce the variance of our parameter estimates. In a Bayesian context, we quantify the information gained from a given dataset by evaluating the using the KL divergence^55^ between the posterior and prior distributions.^40^ In short, evaluates the difference between the updated beliefs once the data has been observed, to the prior beliefs before the data was observed. For a large number of samples, the discrete form of the KL divergence is:

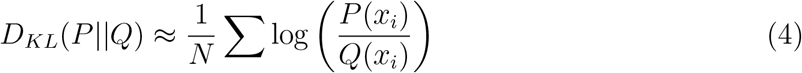

where:

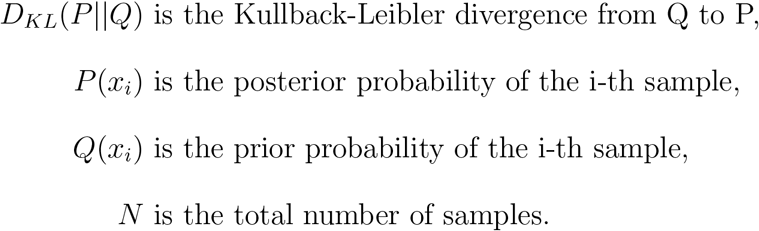

To overcome the issue of low sample number and the ‘noise’ in the posterior distribution, we utilize a Gaussian mixture model (GMM)^56^ to estimate a smooth approximation of the estimated posterior, which are tuned using the Bayesian information criterion (BIC).^57^ This workflow is illustrated in Figure SI 4.

Additionally, we quantify the improved precision of parameter estimates between two posterior distributions using the sum of standard deviations. Specifically, we compare the sum of standard deviations across the marginalized posterior for different datasets:

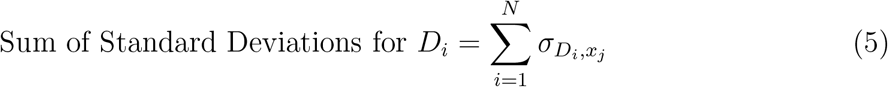

where 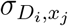 is the standard deviation of the samples for the *x*_*j*_th parameter (N total) and *D*_*i*_th data set.

### Model Selection

In order to compare between possible transporter reaction cycles, we examine the log-likelihoods (3), which in our formulation describe the scaled mean-squared error of the residuals. With Bayesian inference, we generate a distribution of log-likelihoods, which correspond to the log probability of the data given the model parameters. The ability to discern the most likely model depends on the separation of these log-likelihood distributions - if all models are equally likely then their log-likelihood distributions will be overlapping, and if one model is more likely then its log-likelihood distribution will contain the maximum and be separated from the others. Additionally, the model evidence, *P* (*D*) can be estimated using sequential Monte Carlo methods and provides an alternative metric for model comparison^58^ (shown in SI).

### Implementation

We use the Tellurium package^59^ in python^60^ to build human-readable systems biological markup language (SBML)^61^ files using Antimony^62^ and simulate the ODEs using libroad-runner.^63^ For improved reproducibility, we use a .yaml^64^ configuration file to specify the relevant model and data files, as well as the simulated assay conditions and model calibration settings. Bayesian inference with preconditioned Monte Carlo is done using the pocoMC package,^42^ affine invariant ensemble sampling using the emcee package^43^ and maximum likelihood estimation is done using the Scipy Optimize package.^65^ Graphs are generated using Matplotlib,^66^ and numerics are done using numpy.^67^ For Bayesian inference and MLE, multiple replicas are used to check for convergence. The code is available on GitHub: github.com*/*ZuckermanLab*/*Bayesian Transporter, with data available upon request.

## Acknowledgement

The authors thank Katherine Henzler-Wildman, Nathan E. Thomas, and Merissa Brousseau for helpful discussions of SSME experiments; James Faeder, Jeremy Goecks, Peter Jacobs, Herbert Sauro, and Charles Springer for advice regarding the research and manuscript; Michael Grabe and John Rosenberg for valuable discussions of biological transport; and Jeremy Copperman, Aidan Estelle, Sagar Kania, and John Russo for help and advice with the project. This work was supported by NSF grant MCB-2119837.

## Supporting Information Available

Supplemental Information 1: Appendix (.pdf)

## Notes

### Competing Interest Statement

The authors have declared no competing interest.

