## Supplemental Information for "From average transient transporter currents to microscopic mechanism – A Bayesian analysis"

### Supplemental Information (SI): Appendix

August George, Daniel M. Zuckerman
Oregon Health and Science University

#### 1 Transporter Models

In the following section we detail the transporter model differential equations, reaction rate formulation, and considerations for the membrane voltage.

##### 1.1 Model Differential Equations

Here we describe the differential equations governing the four 1:1 antiporter reaction cycles used in the manuscript.

*Cycle 1 differential equations:*

$$
\begin{aligned}
\frac{d[H_{\text{in}}]}{dt} &= k_4^f[IF\_Hb\_Sb] - k_4^r[IF\_Sb][H_{\text{in}}] \\
\frac{d[S_{\text{in}}]}{dt} &= -(k_3^f[IF\_Hb][S_{\text{in}}] - k_3^r[IF\_Hb\_Sb]) \\
\frac{d[OF]}{dt} &= -(k_1^f[OF][H_{\text{out}}] - k_1^r[OF\_Hb]) + k_6^f[OF\_Sb] - k_6^r[OF][S_{\text{out}}] \\
\frac{d[OF\_Hb]}{dt} &= k_1^f[OF][H_{\text{out}}] - k_1^r[OF\_Hb] - (k_2^f[OF\_Hb] - k_2^r[IF\_Hb]) \\
\frac{d[IF\_Hb]}{dt} &= k_2^f[OF\_Hb] - k_2^r[IF\_Hb] - (k_3^f[IF\_Hb][S_{\text{in}}] - k_3^r[IF\_Hb\_Sb]) \\
\frac{d[IF\_Hb\_Sb]}{dt} &= k_3^f[IF\_Hb][S_{\text{in}}] - k_3^r[IF\_Hb\_Sb] - (k_4^f[IF\_Hb\_Sb] - k_4^r[IF\_Sb][H_{\text{in}}]) \\
\frac{d[IF\_Sb]}{dt} &= k_4^f[IF\_Hb\_Sb] - k_4^r[IF\_Sb][H_{\text{in}}] - (k_5^f[IF\_Sb] - k_5^r[OF\_Sb]) \\
\frac{d[OF\_Sb]}{dt} &= k_5^f[IF\_Sb] - k_5^r[OF\_Sb] - (k_6^f[OF\_Sb] - k_6^r[OF][S_{\text{out}}])
\end{aligned}
$$

*Cycle 2 differential equations:*

$$
\begin{aligned}
\frac{d[H_{\text{in}}]}{dt} &= k_4^f[IF\_Hb\_Sb] - k_4^r[IF\_Sb][H_{\text{in}}] \\
\frac{d[S_{\text{in}}]}{dt} &= -(k_3^f[IF\_Hb][S_{\text{in}}] - k_3^r[IF\_Hb\_Sb]) \\
\frac{d[OF\_Hb\_Sb]}{dt} &= -(k_1^f[OF\_Hb\_Sb] - k_1^r[OF\_Hb][S_{\text{out}}]) + k_6^f[OF\_Sb][H_{\text{out}}] - k_6^r[OF\_Hb\_Sb] \\
\frac{d[OF\_Hb]}{dt} &= k_1^f[OF\_Hb\_Sb] - k_1^r[OF\_Hb][S_{\text{out}}] - (k_2^f[OF\_Hb] - k_2^r[IF\_Hb]) \\
\frac{d[IF\_Hb]}{dt} &= k_2^f[OF\_Hb] - k_2^r[IF\_Hb] - (k_3^f[IF\_Hb][S_{\text{in}}] - k_3^r[IF\_Hb\_Sb]) \\
\frac{d[IF\_Hb\_Sb]}{dt} &= k_3^f[IF\_Hb][S_{\text{in}}] - k_3^r[IF\_Hb\_Sb] - (k_4^f[IF\_Hb\_Sb] - k_4^r[IF\_Sb][H_{\text{in}}]) \\
\frac{d[IF\_Sb]}{dt} &= k_4^f[IF\_Hb\_Sb] - k_4^r[IF\_Sb][H_{\text{in}}] - (k_5^f[IF\_Sb] - k_5^r[OF\_Sb]) \\
\frac{d[OF\_Sb]}{dt} &= k_5^f[IF\_Sb] - k_5^r[OF\_Sb] - (k_6^f[OF\_Sb][H_{\text{out}}] - k_6^r[OF\_Hb\_Sb])
\end{aligned}
$$

*Cycle 3 differential equations:*

$$\frac{d[H_{\text{in}}]}{dt} = k_3^f[IF\_Hb] - k_3^r[IF][H_{\text{in}}]$$
$$\frac{d[S_{\text{in}}]}{dt} = -(k_4^f[IF][S_{\text{in}}] - k_4^r[IF\_Sb])$$
$$\frac{d[OF]}{dt} = -(k_1^f[OF][H_{\text{out}}] - k_1^r[OF\_Hb]) + k_6^f[OF\_Sb] - k_6^r[OF][S_{\text{out}}]$$
$$\frac{d[OF\_Hb]}{dt} = k_1^f[OF][H_{\text{out}}] - k_1^r[OF\_Hb] - (k_2^f[OF\_Hb] - k_2^r[IF\_Hb])$$
$$\frac{d[IF\_Hb]}{dt} = k_2^f[OF\_Hb] - k_2^r[IF\_Hb] - (k_3^f[IF\_Hb] - k_3^r[IF][H_{\text{in}}])$$
$$\frac{d[IF]}{dt} = k_3^f[IF\_Hb] - k_3^r[IF][H_{\text{in}}] - (k_4^f[IF][S_{\text{in}}] - k_4^r[IF\_Sb])$$
$$\frac{d[IF\_Sb]}{dt} = k_4^f[IF][S_{\text{in}}] - k_4^r[IF\_Sb] - (k_5^f[IF\_Sb] - k_5^r[OF\_Sb])$$
$$\frac{d[OF\_Sb]}{dt} = k_5^f[IF\_Sb] - k_5^r[OF\_Sb] - (k_6^f[OF\_Sb] - k_6^r[OF][S_{\text{out}}])$$

*Cycle 4 differential equations:*

$$\frac{d[H_{\text{in}}]}{dt} = k_3^f[IF\_Hb] - k_3^r[IF][H_{\text{in}}]$$
$$\frac{d[S_{\text{in}}]}{dt} = -(k_4^f[IF][S_{\text{in}}] - k_4^r[IF\_Sb])$$
$$\frac{d[OF\_Hb\_Sb]}{dt} = -(k_1^f[OF\_Hb\_Sb] - k_1^r[OF\_Hb][S_{\text{out}}]) + k_6^f[OF\_Sb][H_{\text{out}}] - k_6^r[OF\_Hb\_Sb]$$
$$\frac{d[OF\_Hb]}{dt} = k_1^f[OF\_Hb\_Sb] - k_1^r[OF\_Hb][S_{\text{out}}] - (k_2^f[OF\_Hb] - k_2^r[IF\_Hb])$$
$$\frac{d[IF\_Hb]}{dt} = k_2^f[OF\_Hb] - k_2^r[IF\_Hb] - (k_3^f[IF\_Hb] - k_3^r[IF][H_{\text{in}}])$$
$$\frac{d[IF]}{dt} = k_3^f[IF\_Hb] - k_3^r[IF][H_{\text{in}}] - (k_4^f[IF][S_{\text{in}}] - k_4^r[IF\_Sb])$$
$$\frac{d[IF\_Sb]}{dt} = k_4^f[IF][S_{\text{in}}] - k_4^r[IF\_Sb] - (k_5^f[IF\_Sb] - k_5^r[OF\_Sb])$$
$$\frac{d[OF\_Sb]}{dt} = k_5^f[IF\_Sb] - k_5^r[OF\_Sb] - (k_6^f[OF\_Sb][H_{\text{out}}] - k_6^r[OF\_Hb\_Sb])$$

In these equations, [X] denotes the concentration of reaction state 'X', OF and IF represent the outward-facing and inward-facing conformations. The transported ion (H) and substrate (S) are compartmentalized into either outside the liposome ($X_{out}$), inside the liposome ($X_{in}$), or bound to the transporter protein ($X_b$). The reaction rate constant $k_i^f$ corresponds to the rate constant for reaction i, in the forward clockwise direction 'f', with the counterclockwise direction denoted with 'r'. We note that for the simulated SSME experiment, the $H_{\text{out}}$ and $S_{\text{out}}$ concentrations remain fixed during each assay stage.

#### 1.2 Reaction Rate Constants

To account for the effects of the membrane voltage on the transport of charged chemical species we use a modified Arrhenius rate formulation using Eyring rate theory [4, 8]:

$$k_i^j(V) = k_i^j(0)\exp(-\epsilon_i^j VF/(RT)) \tag{1}$$

where $k_i^j(0)$ is the rate constant at zero voltage, F is Faraday's constant, R is the gas constant, T is the absolute temperature, V is the membrane voltage, and $\epsilon_i^j$ is the fractional charge transported during reaction step $i$, in the $j$ (forward or reverse) direction. In our model, it is assumed that all the charge is transported when an ion, $H^+$, binds or unbinds inside the liposome. For the cycle 1 and 2 models, $\epsilon_4^f = 1$, $\epsilon_4^r = -1$, and all other $\epsilon = 0$. For the cycle 3 and 4 models, $\epsilon_3^f = 1$, $\epsilon_3^r = -1$, and all other $\epsilon = 0$.

The membrane voltage, V, is time-dependent, and we approximate the complex dynamics[11, 5, 22] using an idealized capacitor model:

$$V_m(t) = Q(t)/C_m = \int I(t)dt/C_m \tag{2}$$

where $I(t)$ is the current of the transported ion, derived from the change in concentration of the ion inside the liposome, and $C_m$ is the capacitance of the liposome membrane, which is known empirically. For this study we assume a specific membrane capacitance of 0.5e-6 $\frac{F}{cm^2}$ [6], and an approximate average surface area of 2e-10 $cm^2$ [27], which yields a membrane capacitance ($C_m$) of 1e-16 F. Empirical results for our model suggested a negligible effect of the membrane voltage on the rate constants and observables (see below).

At equilibrium, the products of the forward reaction rate constants must equal the products of the reverse reaction rate constants for each step to preserve microscopic reversibility [10, 3]. This necessarily means that one rate constant is not independent. In our implementation, we explicitly set $k_6^-$ as the non-independent rate constant defined by the ratio of remaining forward and reverse reaction rate constants.

#### 1.3 Effects of membrane potential on rate constants

We examine the relationship between membrane voltage, charge, and current for our transporter model and simulated SSME assays: $V_m(t) = Q(t)/C_m = \int I(t)dt/C_m$

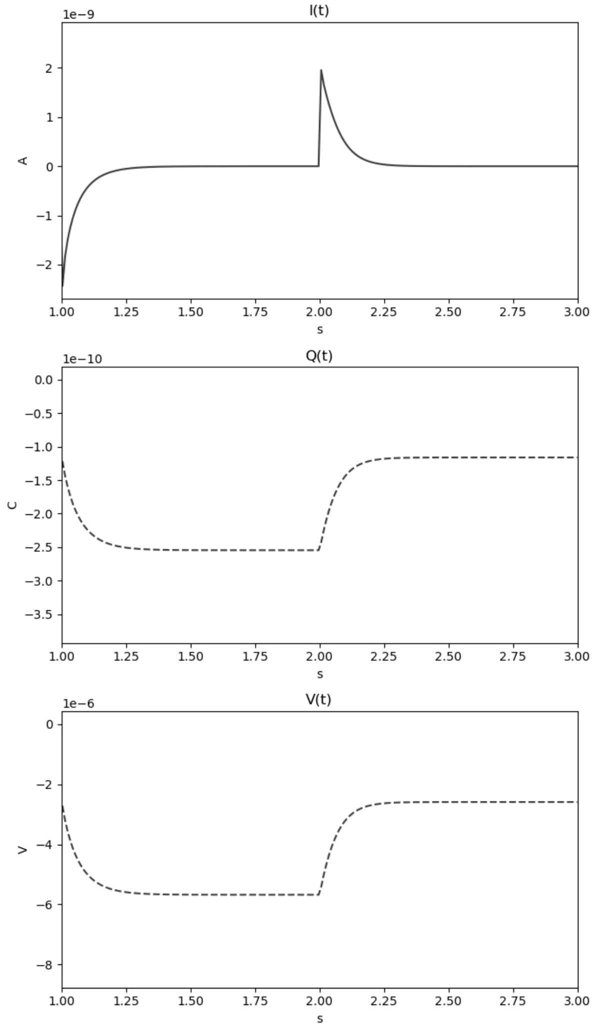


Figure 1: **Characterizing current, charge, and membrane voltage.** The total current, transported charge, and membrane voltage for a simulated SSME assay for t=1-3 seconds. The equilibration phase creates a buildup of positive charge on the membrane's extracellular side, giving a negative charge Q(t) by convention.

We find that the membrane voltage has a negligible effect on the reaction rate constants and the resulting current, for the conditions studied. Consider the modified rate constant form: $k(V) = k_0 * \exp(-\epsilon * \frac{VF}{RT})$. While exact values will vary depending on the transporter and experimental setup, the peak of the net currents is often in the nA range and transport a total charge in the nC range [1, 2],. This results in a per liposome membrane total charge of 1e-23 C, which when divided by a typical liposome membrane capacitance of 1e-16 F results in a voltage in the $\mu$V range. If we assume a maximum voltage of 1e-5 V, room temperature of 298K, and $\epsilon = 1$, this yields: $k(V) \approx k_0 * 0.9996$. This suggests that the membrane voltage adjusts the rate constants by a negligible amount for the conditions studied. We note that under physiological conditions, the resting membrane voltage is typically maintained in the 50-100mV magnitude range. Under these conditions, the membrane voltage will significantly adjust the rate constants, with $k(V) \approx k_0 * 0.15$ for 50 mV.

We compare the Bayesian inference results on the same transporter model using a dynamic voltage, $k(V) = k_0 * \exp(-\epsilon * \frac{VF}{RT})$, and under a zero voltage condition, $k(V) = k_0$. As expected from our model described above, we find a negligible difference between the two voltage formulations. Further work will investigate the membrane potential in greater detail.

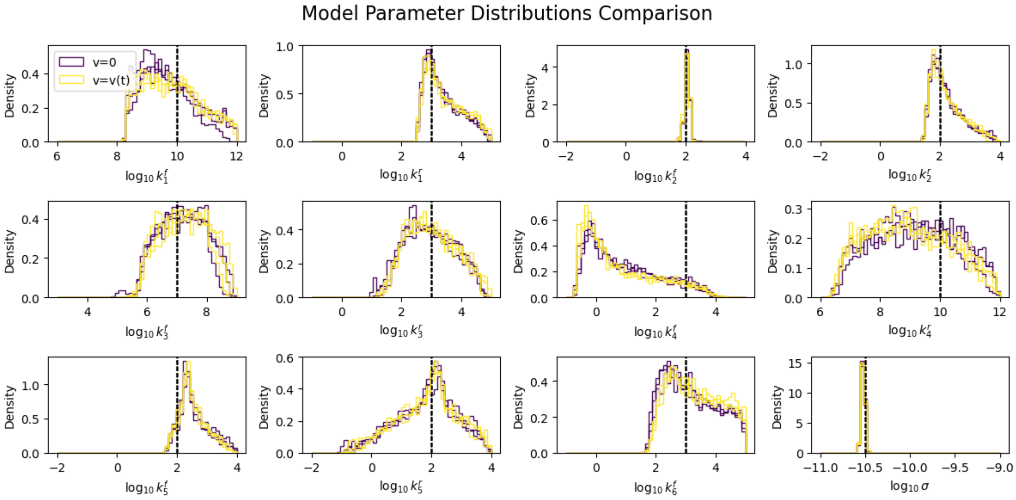


Figure 2: **Parameter distributions.** The 1D parameter distributions with and without dynamic voltage. Both voltage formulations result in nearly identical parameter distributions, with three replicas for each formulation shown. The ground truth values are shown in vertical dashed lines for reference.

### 2 Synthetic SSME assay conditions

We assume uniformity of the liposomes, such that the concentrations are the same between each liposome, and single liposome volumes are equal to the total volume divided by the number of liposomes. We use parameter values motivated by SSME experiments and simulations done for Gdx and EmrE proteins [27, 12, 29].

We only explicitly model the driving ion and substrate that are directly coupled to the transport process studied under SSME-like conditions. Therefore we do not explicitly model water, anions, or other chemical species used in the bath solution. We assume that these species concentrations are controlled such that they have a negligible effect on transport during an SSME assay, and that their effects are accounted for by the rate constant values.

More specifically, we base our approach on related SSME experiments [29] for the Gdx protein, which holds the chloride ($Cl^-$) anion concentrations fixed throughout the bath solution via a controlled buffer, removing potential confounding effects from varied anion concentrations.

The differential equations are integrated with the CVODES integrator with an absolute tolerance of 1e-15, relative tolerance of 1e-12. A separate integration run is used for each of the stages (i.e. t=0-1s, t=1-2s, t=2-3s, etc.) to account for the discrete-time events. For the purposes of calculating the log-likelihood, we omit the data from the equilibration stage (t=0 to t=1 s), and for the remaining assay stages, we remove the first data point and the last one-hundred data points (i.e., after a steady-state is reached).

The net current is calculated from the change in internal ion concentrations of a single liposome – converting from the change in molar concentration to current, and multiplying by the number of liposomes in the experiment.

$$I_{\text{net}}(t) = I_{\text{liposome}}(t) N_{\text{liposomes}} = (\frac{d[H_{in}^+]}{dt} \text{Vol}_{\text{in}} N_{Av} z) N_{\text{liposomes}} \quad (3)$$

where $\text{Vol}_{\text{in}}$ is the internal volume of a single liposome embedded with transporters, $N_{Av}$ is Avogadro's constant, $z$ is the elementary charge of an $H^+$ ion, and $N_{\text{liposomes}}$ is the total number of liposomes in the SSME assay. Here we assume uniformity across the aggregate of liposomes.

| | |
|---|---|
| N liposomes total | 1e11 |
| N transporters total | 5e12 |
| Total external volume | 70 uL |
| Total internal volume | 0.07 uL |
| Total membrane volume | 0.02 uL |
| Liposome membrane capacitance | 2.4e-16 F |

Table 1: Summary of parameters

#### 2.1 Transporter model state definitions and initial concentrations

We use the following definitions for biochemical states and species:

| | |
|---|---|
| OF | Outward-facing conformation |
| OF_Sb | Outward-facing conformation with substrate bound |
| OF_Hb | Outward-facing conformation with ion bound |
| OF_Hb_Sb | Outward-facing conformation with ion and substrate bound |
| IF_Sb | Inward-facing conformation with substrate bound |
| IF_Hb | Inward-facing conformation with ion bound |
| IF_Hb_Sb | Inward-facing conformation with ion and substrate bound |
| $H_{out}$ | External (i.e. bath) ion |
| $S_{out}$ | External (i.e. bath) substrate |
| $H_{in}$ | Internal (i.e. inside lissome) ion |
| $S_{in}$ | Internal (i.e. inside lissome) substrate |

We use the following initial concentrations for each model:

Table 2: Model 1: Species Initial Concentrations (M)

| Species Label | Initial Concentration |
|---|---|
| OF | 0.00044 |
| OF_Hb | 0 |
| IF_Hb | 0 |
| IF_Hb_Sb | 0 |
| IF_Sb | 0 |
| OF_Sb | 0 |
| H_in | 1.e-7 |
| S_in | 1.e-3 |
| H_out | 1.e-7 |
| S_out | 1.e-3 |

Table 3: Model 2: Species Initial Concentrations (M)

| Species Label | Initial Concentration |
|---|---|
| OF_Hb_Sb | 0.00044 |
| OF_Hb | 0 |
| IF_Hb | 0 |
| IF_Hb_Sb | 0 |
| IF_Sb | 0 |
| OF_Sb | 0 |
| H_in | 1.e-7 |
| S_in | 1.e-3 |
| H_out | 1.e-7 |
| S_out | 1.e-3 |

Table 4: Model 3: Species Initial Concentrations (M)

| Species Label | Initial Concentration |
|---|---|
| OF | 0.00044 |
| OF_Hb | 0 |
| IF_Hb | 0 |
| IF | 0 |
| IF_Sb | 0 |
| OF_Sb | 0 |
| H_in | 1.e-7 |
| S_in | 1.e-3 |
| H_out | 1.e-7 |
| S_out | 1.e-3 |

Table 5: Model 4: Species Initial Concentrations (M)

| Species Label | Initial Concentration |
|---|---|
| OF_Hb_Sb | 0.00044 |
| OF_Hb | 0 |
| IF_Hb | 0 |
| IF | 0 |
| IF_Sb | 0 |
| OF_Sb | 0 |
| H_in | 1.e-7 |
| S_in | 1.e-3 |
| H_out | 1.e-7 |
| S_out | 1.e-3 |

#### 2.2 Synthetic assays for experiment recommendation

The external concentrations of the ion (H) and substrate (S) are pertubed by different amounts for different assays. Then datasets are generated by combining or repeating the assay conditions. We note that not all datasets are used in this study. The concentration settings for each protocol and visualizations of the data traces are shown below:

| **Protocol** | **Synthetic Data** | $H_{out}$ **Concentration in M** |
|---|---|---|
| 1 | Experiment 1 | 1.e-7, 0.5e-7, 1.e-7 |
| 2 | Experiment 2 | 1.e-7, 0.5e-7, 1.e-7 |
| - | Experiment 3 | 1.e-7, 0.5e-7, 1.e-7 |
| - | Experiment 4 | 1.e-7, 0.5e-7, 1.e-7 |
| 3 | Experiment 1+2 | 1.e-7, 0.5e-7, 1.e-7, 0.5e-7, 1.e-7 |
| 4 | Experiment 1+2+3+4 | 1.e-7, 0.5e-7, 1.e-7, 0.5e-7, 1.e-7, 0.5e-7, 1.e-7, 0.5e-7, 1.e-7 |
| 5 | Experiment 1x4 | 1.e-7, 0.5e-7, 1.e-7, 0.5e-7, 1.e-7, 0.5e-7, 1.e-7, 0.5e-7, 1.e-7 |

Table 6: Buffer concentration sequence for experiment recommendation data sets - $H_{out}$ concentrations. Two $H_{out}$ conditions are studied, 1e-7 and 0.5e-7 M, which correspond to external pHs of 7 and 7.3, respectively.

| **Protocol** | **Synthetic Data** | $S_{out}$ **Concentration in M** |
|---|---|---|
| 1 | Experiment 1 | 1.e-3, 1.0e-3, 1.e-3 |
| 2 | Experiment 2 | 1.e-3, 0.353e-3, 1.e-3 |
| - | Experiment 3 | 1.e-3, 0.5e-3, 1.e-3 |
| - | Experiment 4 | 1.e-3, 0.25e-3, 1.e-3 |
| 3 | Experiment 1+2 | 1.e-3, 1.0e-3, 1.e-3, *0.353e-3*, *1.e-3* |
| 4 | Experiment 1+2+3+4 | 1.e-3, 1.0e-3, 1.e-3, 0.5e-3, 1.e-3, *0.353e-3*, *1.e-3*, 0.25e-3, 1.e-3 |
| 5 | Experiment 1x4 | 1.e-3, 1.0e-3, 1.e-3, 1.0e-3, 1.e-3, 1.0e-3, 1.e-3, 1.0e-3, 1.e-3 |

Table 7: Buffer concentration sequence for experiment recommendation data sets - $S_{out}$ concentrations. We note that in protocol 4, the experiment 2 perturbation is third sequentially.

Finally, we show the synthetic SSME-like data traces resulting from the simulated assay conditions below:

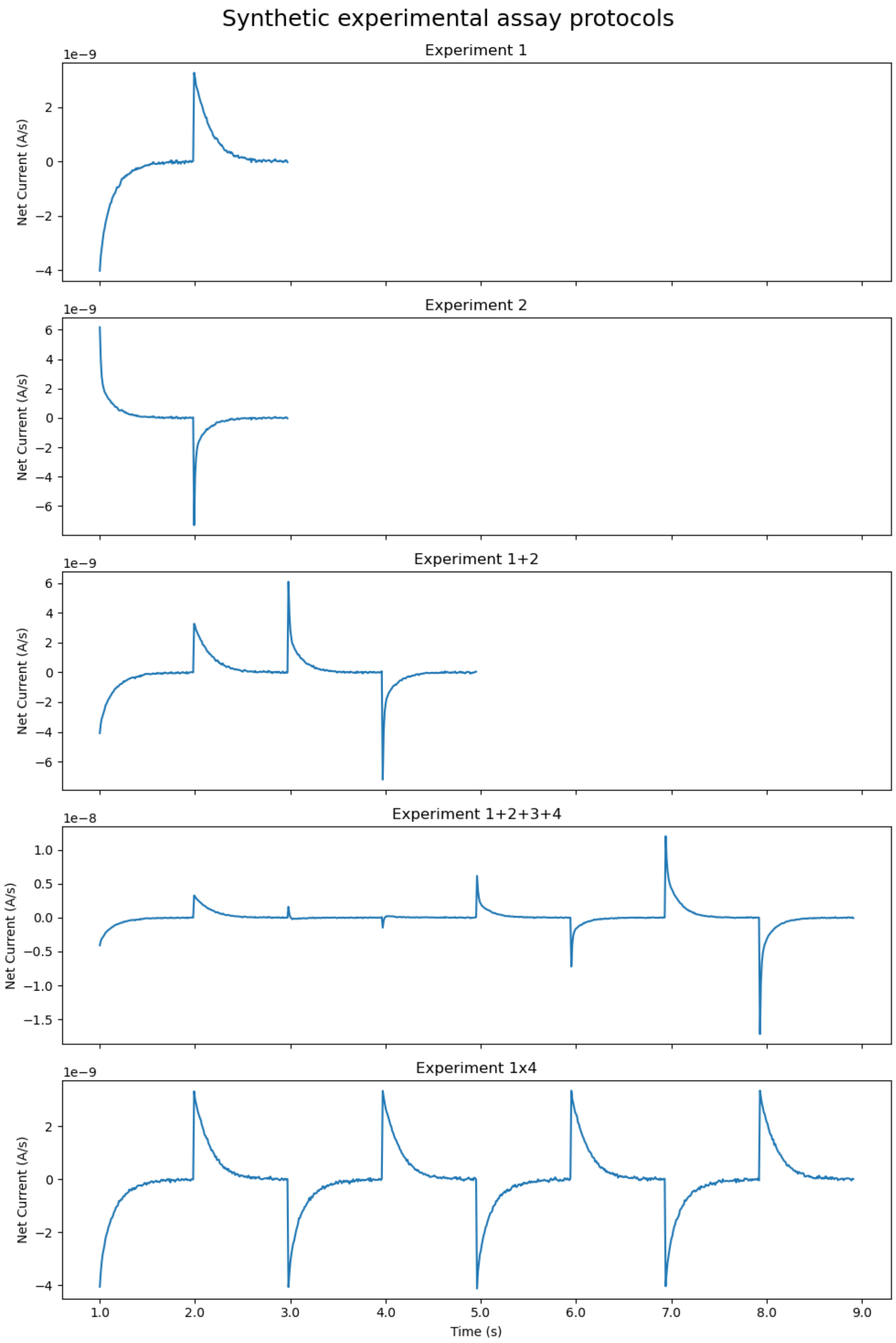


Figure 3: **Synthetic SSME Assay Datasets.** Selected datasets corresponding to different perturbation amounts, combinations of experiments, and technical replicas.

### 3 Probabilistic modeling

In the following section we describe in detail the probabilistic model including the likelihood function, prior ranges, and parameter nominal values.

#### 3.1 Log-likelihood function

As mentioned in the manuscript, we use a Normal log-likelihood distribution for Bayesian inference:

$$L(\theta, \sigma^2|D) = -\frac{n}{2}ln(2\pi) - \frac{n}{2}ln(\sigma^2) - \frac{1}{2\sigma^2}\sum_{i=1}^{n}(D_i - D_{pred}(\theta)_i)^2 \quad (4)$$

where the predicted data, $D_{pred}$, is generated from equation 3, $\sigma^2$ is the (unknown) variance, and $D$ is the given observed (synthetic) data.

#### 3.2 Rate constant priors

We use extremely broad uniform priors covering six orders of magnitude, with a log10 transformation for the reaction rate constants. Below the log10 rate constant ranges and synthetic ground truth (nominal) values are shown for each transport cycle model:

Table 8: Model1: Priors

| Name | Bounds | Nominal |
|---|---|---|
| log10_k1_f | [6, 12] | 10 |
| log10_k1_r | [-1, 5] | 3 |
| log10_k2_f | [-2, 4] | 2 |
| log10_k2_r | [-2, 4] | 2 |
| log10_k3_f | [3, 9] | 7 |
| log10_k3_r | [-1, 5] | 3 |
| log10_k4_f | [-1, 5] | 3 |
| log10_k4_r | [6, 12] | 10 |
| log10_k5_f | [-2, 4] | 2 |
| log10_k5_r | [-2, 4] | 2 |
| log10_k6_f | [-1, 5] | 3 |

Table 9: Model 2: Priors

| Name | Bounds | Nominal |
|---|---|---|
| log10_k1_f | [-1, 5] | 3 |
| log10_k1_r | [3, 9] | 7 |
| log10_k2_f | [-2, 4] | 2 |
| log10_k2_r | [-2, 4] | 2 |
| log10_k3_f | [3, 9] | 7 |
| log10_k3_r | [-1, 5] | 3 |
| log10_k4_f | [-1, 5] | 3 |
| log10_k4_r | [6, 12] | 10 |
| log10_k5_f | [-2, 4] | 2 |
| log10_k5_r | [-2, 4] | 2 |
| log10_k6_f | [6, 12] | 10 |

Table 10: Model 3: Priors

| Name | Bounds | Nominal |
|---|---|---|
| log10_k1_f | [6, 12] | 10 |
| log10_k1_r | [-1, 5] | 3 |
| log10_k2_f | [-2, 4] | 2 |
| log10_k2_r | [-2, 4] | 2 |
| log10_k3_f | [-1, 5] | 3 |
| log10_k3_r | [6, 12] | 10 |
| log10_k4_f | [3, 9] | 7 |
| log10_k4_r | [-1, 5] | 3 |
| log10_k5_f | [-2, 4] | 2 |
| log10_k5_r | [-2, 4] | 2 |
| log10_k6_f | [-1, 5] | 3 |

Table 11: Model 4: Priors

| Name | Bounds | Nominal |
|---|---|---|
| log10_k1_f | [-1, 5] | 3 |
| log10_k1_r | [3, 9] | 7 |
| log10_k2_f | [-2, 4] | 2 |
| log10_k2_r | [-2, 4] | 2 |
| log10_k3_f | [-1, 5] | 3 |
| log10_k3_r | [6, 12] | 10 |
| log10_k4_f | [3, 9] | 7 |
| log10_k4_r | [-1, 5] | 3 |
| log10_k5_f | [-2, 4] | 2 |
| log10_k5_r | [-2, 4] | 2 |
| log10_k6_f | [6, 12] | 10 |

#### 3.3 Nuisance parameters and priors

We assume that the residual errors in the SSME data are from a Normal distribution with zero mean and unknown variance:

$$D_{obs} = D_{true} + \delta$$

where $\delta \sim \text{Normal}(0, \sigma^2)$

We introduce additional sources of uncertainty by using scaling factors in the initial transporter (e.g. protein) concentrations, the external bath perturbation concentrations, and the observed data.

$$\begin{aligned} OF(0)_{obs} &= f_{transporter} \cdot OF(0)_{true} \\ OF\_Hb\_Sb(0)_{obs} &= f_{transporter} \cdot OF\_Hb\_Sb(0)_{true} \\ H_{out_{obs}} &= f_{H_{out}} \cdot H_{out_{true}} \\ S_{out_{obs}} &= f_{S_{out}} \cdot S_{out_{true}} \\ D_{obs} &= f_{bias} \cdot D_{true} + \delta \end{aligned}$$

We use uniform priors for the nuisance parameters, with a log10 scale for the standard deviation.

| nuisance parameter | nominal value | prior range |
|---|---|---|
| log10 $\sigma$ | -10.5 | [-11,-9] |
| $f_{\text{transporter}}$ | 1 | [0.8,1.2] |
| $f_{H_{out}}$ | 1 | [0.8,1.2] |
| $f_{S_{out}}$ | 1 | [0.8,1.2] |
| $f_{\text{bias}}$ | 1 | [0.8,1.2] |

Table 12: Priors used for nuisance parameters

### 4 Information Quantification

As mentioned in the manuscript, we use Gaussian mixture models [25] to generate a smooth approximation of the posterior from the Bayesian samples. GMMs are a probabilistic model that aim to fit a collection of weighted multidimensional Gaussians to the target (in this case posterior) distribution. They generate a smooth analytical function that can easily generate a large number of samples, or be used directly for calculations.

$$p(\mathbf{x}) = \sum_{i=1}^{K} \pi_i \mathcal{N}(\mathbf{x}|\boldsymbol{\mu}_i, \boldsymbol{\Sigma}_i) \tag{5}$$

where:

- $\mathbf{x}$ is the data vector.
- $\pi_i$ is the mixture weight for the $i$-th Gaussian, and all weights sum to 1.
- $\mathcal{N}(\mathbf{x}|\boldsymbol{\mu}_i, \boldsymbol{\Sigma}_i)$ is the $i$-th Gaussian distribution with mean $\boldsymbol{\mu}_i$ and covariance $\boldsymbol{\Sigma}_i$.
- $K$ is the total number of Gaussian distributions in the mixture model.

A key hyper-parameter governing GMMs is the number of Gaussians, K. We use an elbow plot method [16] to select K. Here we iterate from small to larger K values, calculate an information criteria value that balances the goodness of fit against the number of parameters, and stop when a minimum has been reached.

For this study, we used the Bayesian Information Criterion (BIC)[20]:

$$BIC = \ln(n)k - 2\ln(\hat{L}) \tag{6}$$

where $n$ is the number of observations, $\hat{L}$ is the computed likelihood and $k$ is the number of free parameters in the model. This entire process of information quantification is outlined in figure 4 below.

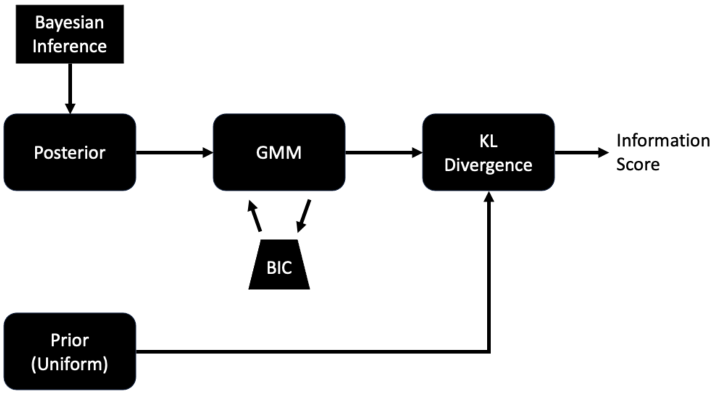


Figure 4: **Information Quantification Workflow.** Bayesian inference generates an estimated posterior which is approximated using a Gaussian mixture model (GMM). The number of Gaussians is optimized using the Bayesian information criterion that penalizes over-fitting. The GMM, representing the posterior, can then calculate the KL divergence with the prior distribution.

### 5 Additional Marginal Posterior Distributions

For completeness, we show the marginal posterior distributions for the cycle 1 experiment 2, and cycles 2, 3, and 4 using the experiment 1 and combined experiments 1-4 datasets. We note that for all the plots below, the referece values (from cycle 1) are shown with a vertical dashed line.

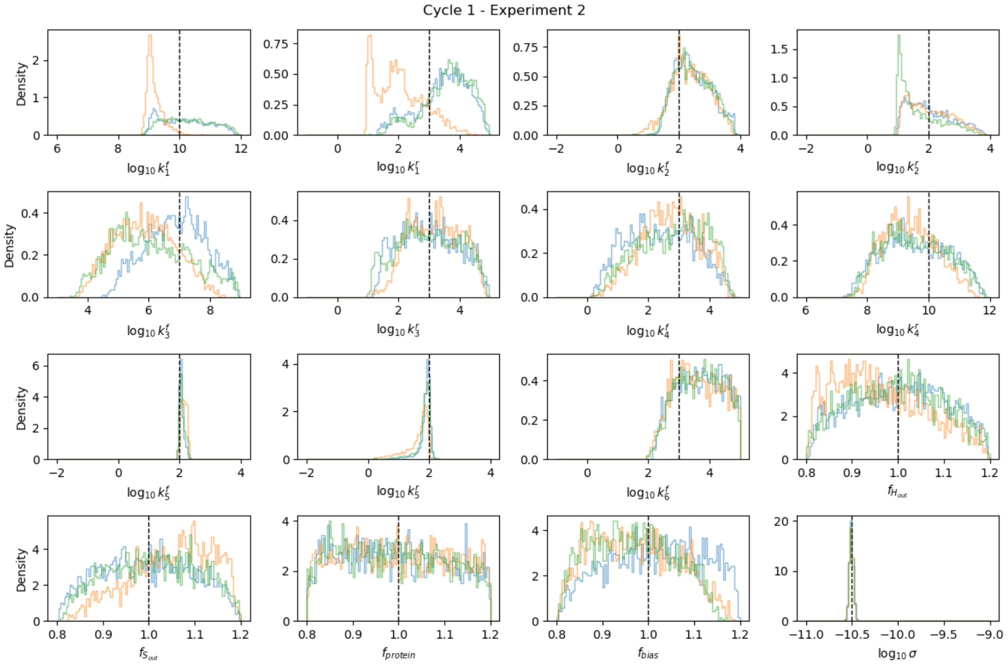


Figure 5: **Posterior for cycle 1 experiment 2 with replicas.**

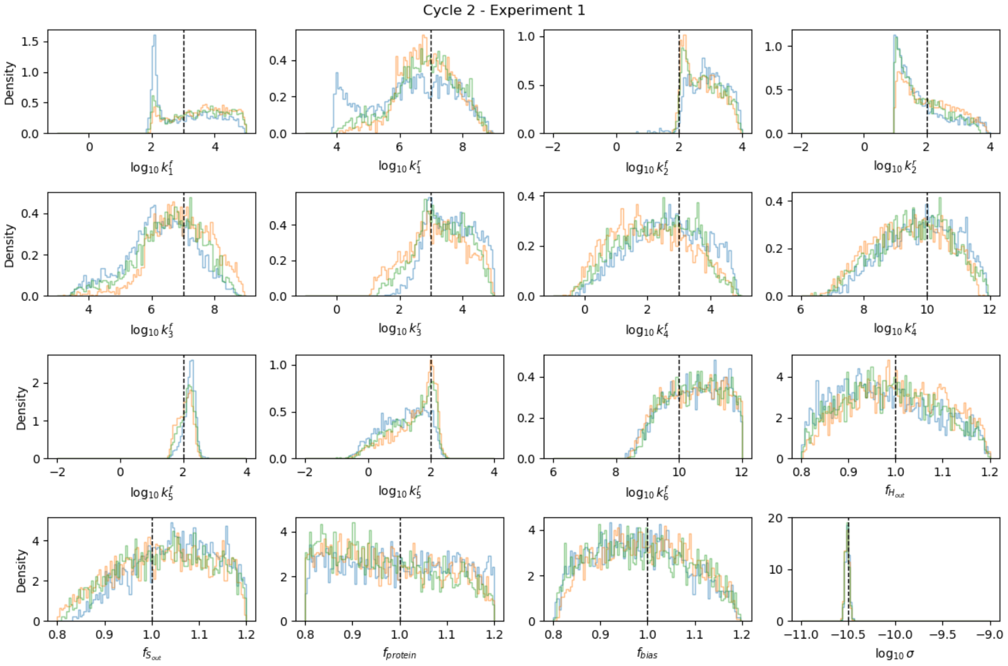


Figure 6: **Posterior for cycle 2 experiment 1 with replicas.**

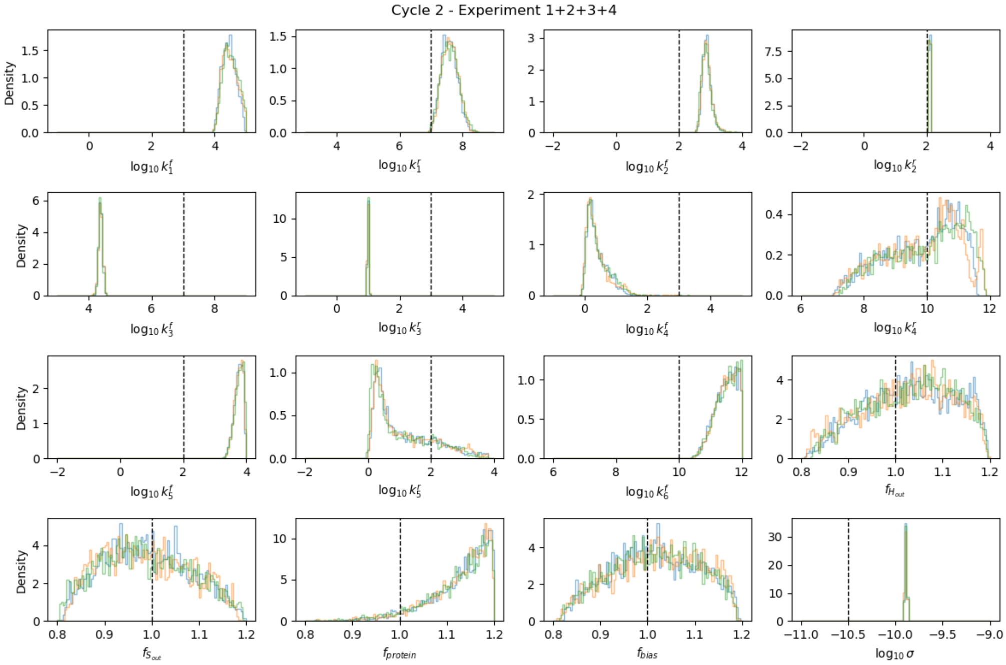


Figure 7: **Posterior for cycle 2 experiment 1+2+3+4 with replicas.**

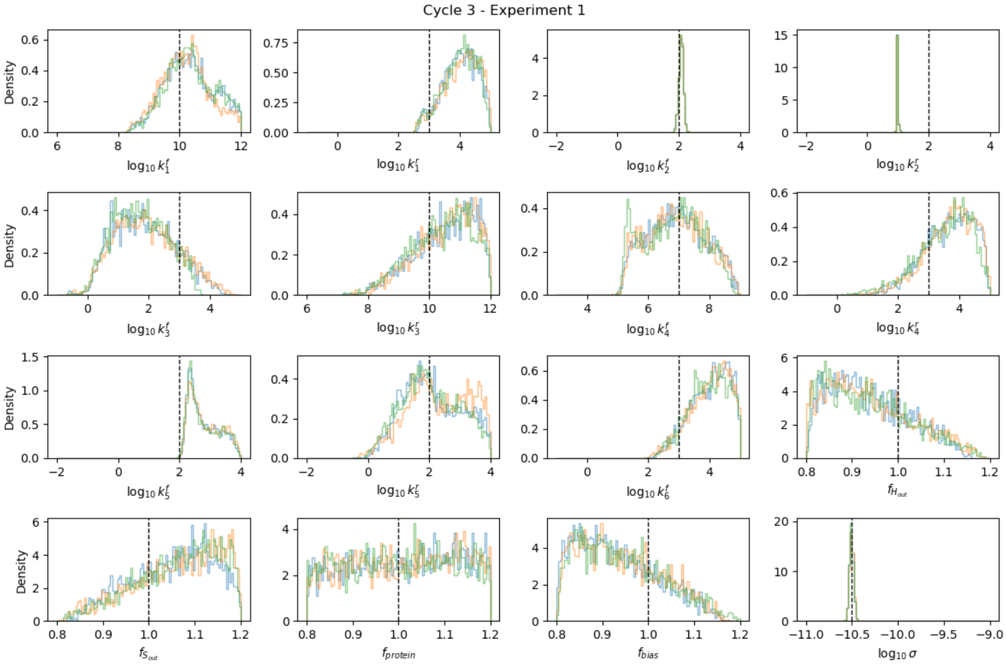


Figure 8: **Posterior for cycle 3 experiment 1 with replicas.**

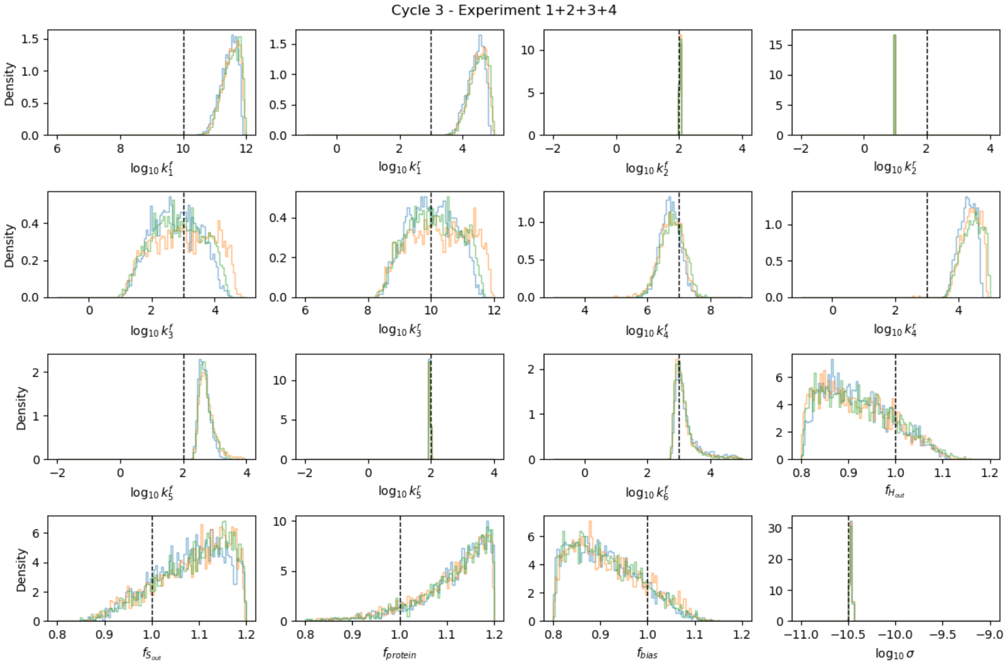


Figure 9: **Posterior for cycle 3 experiment 1+2+3+4 with replicas.**

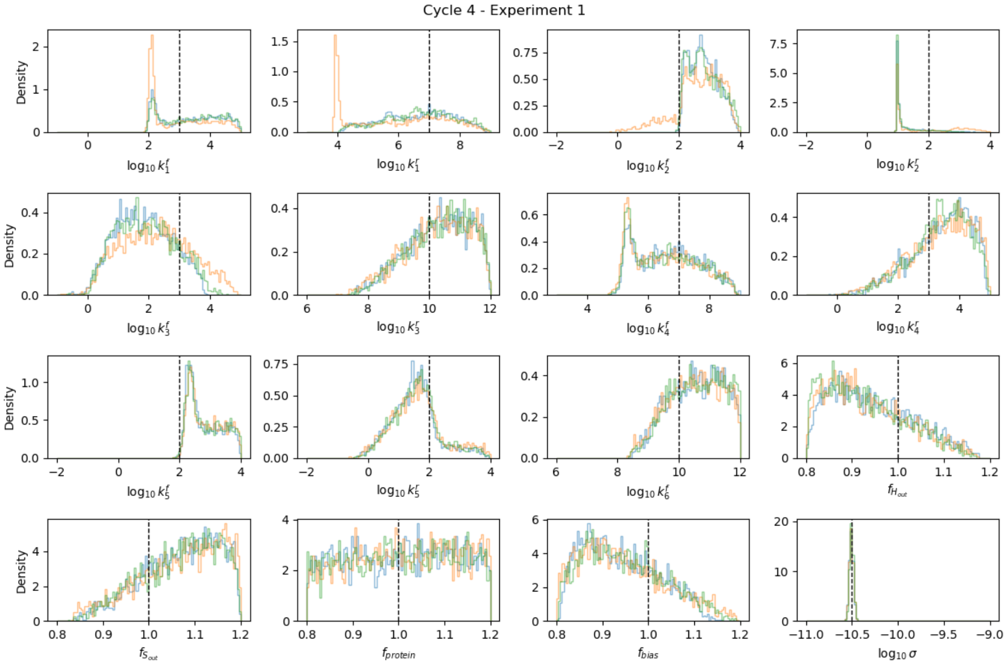


Figure 10: **Posterior for cycle 4 experiment 1 with replicas.**

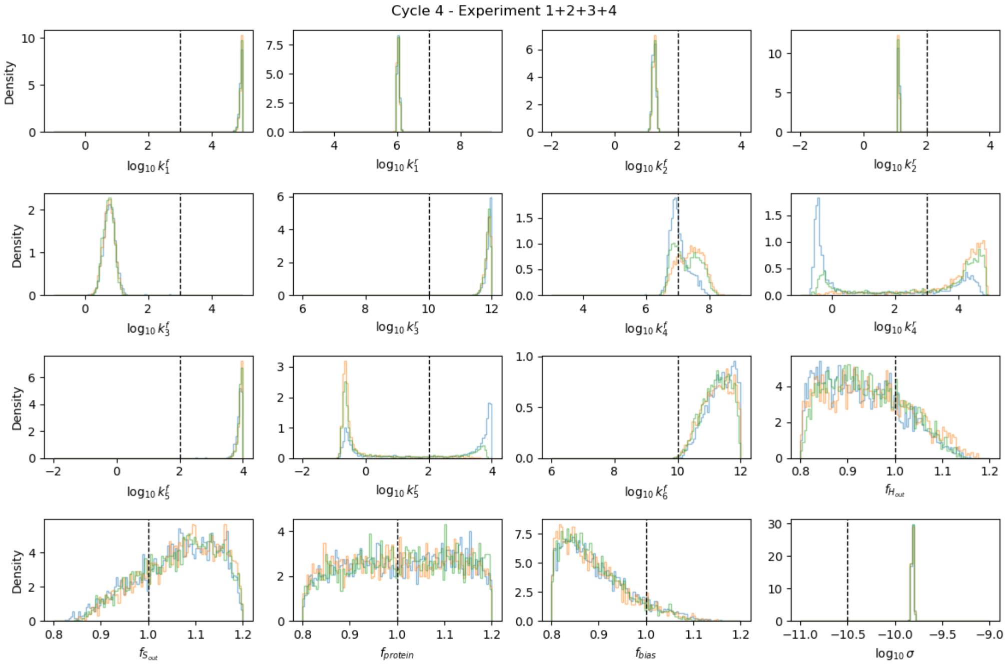


Figure 11: **Posterior for cycle 4 experiment 1+2+3+4 with replicas.**

Finally, we show the corner plot for the combined dataset of experiments 1, 2, 3 and 4 using the cycle 1 model. Here the data from three replica runs are concatenated and the 1D and 2D marginal posteriors are plotted. We see that certain pairs of rate constants are correlated, which suggests that ratios of these rate constants pairs (i.e. $K_D$ values) are identifiable.

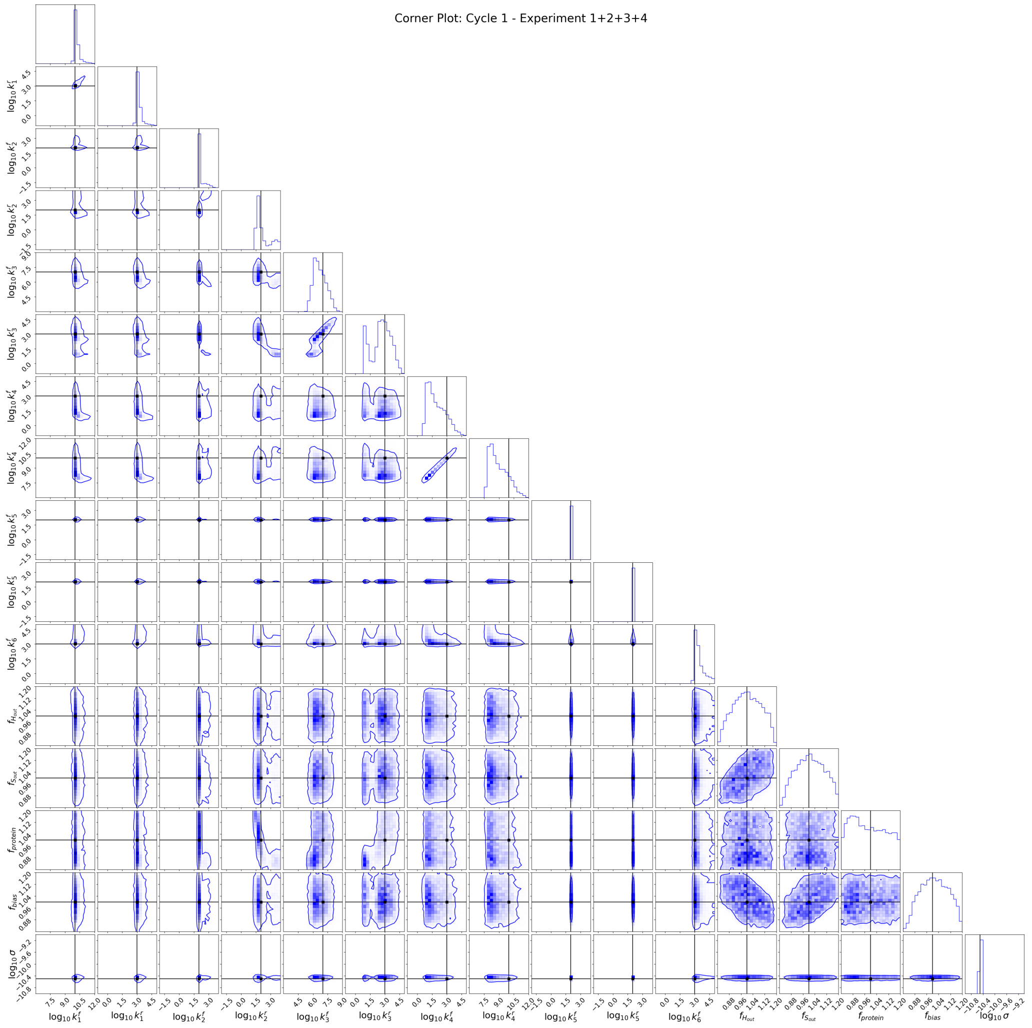


Figure 12: **Corner plot for the combined dataset of experiments 1, 2, 3, and 4 for cycle 1.** The reference values are shown in black, with 95% credible interval shown as a contour. We note that log10 $k_3^f$ and $k_3^r$ as well as $k_4^f$ and $k_4^r$ rate constant pairs are correlated, which suggests that their ratios (i.e. $K_D$ values) are identifiable.

### 6 Comparison of 16D and 12D models

Below we present the sum of standard deviations for the 16D and 12D models.

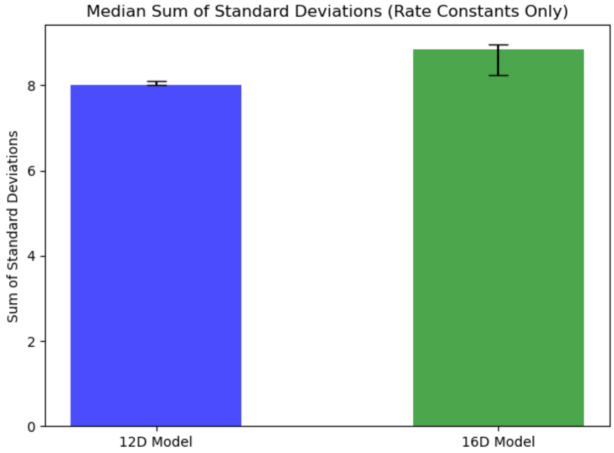


Figure 13: **The median sum of standard deviations for the 16D and 12D model.** As expected the 12D model has a lower sum of standard deviations. The minimum and maximum range is also shown for reference.

### 7 Model Selection using Model Evidence

Sequential Monte Carlo methods (like the pocoMC implementation of preconditioned Monte Carlo) can estimate the normalization constant in Bayes' Theorem, P(D), referred to as the model evidence. Comparing the difference (or ratio) of these values across different models can aid with model identification and selection. Below we show the model evidence for all four cycles using both a relatively uniformative and informative dataset (i.e., Experiment 1 vs Experiment 1+2+3+4, respectively). We find that for the uninformative dataset the differences in the log-evidences are much less significant than with the more informative dataset - supporting our results in the manuscript.

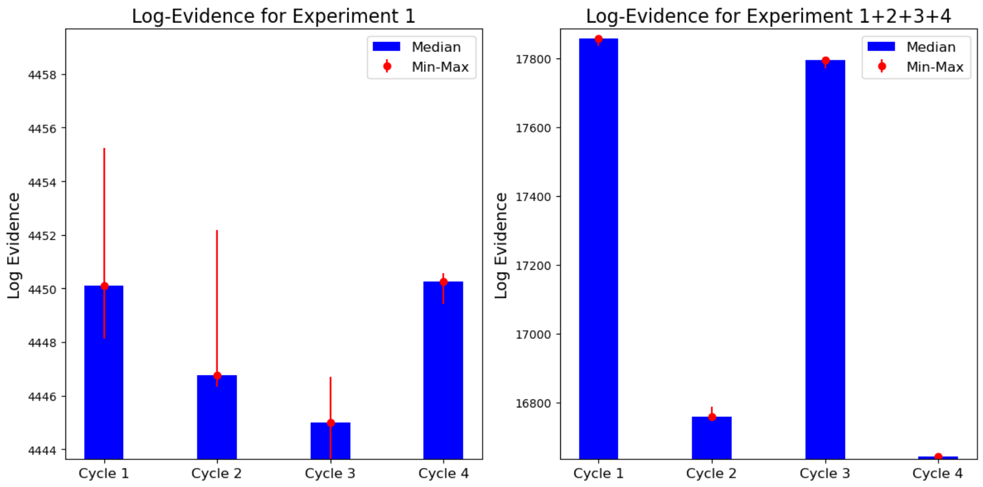


Figure 14: **Comparison of Log-Evidence.** The median log-evidence estimated across multiple replicas for an uninformative (left) and informative (right) dataset with minimum and maximum range shown in red. We find that the relative differences between the cycle 1 and other cycle log-evidences are significant for the informative dataset - suggesting that cycle 1 is the most likely candidate model.

### 8 Comparison of Bayesian and MLE Algorithms

To validate our pipeline using preconditioned Monte Carlo (PMC) [13] we examine several different parameter estimation strategies for comparison. For Bayesian inference we use the affine invariant ensemble sampler (AIES) [7] as an alternative method which uses multiple MCMC chains (i.e. walkers). In AIES, each walker samples the posterior space via correlated jumps based on the position of other walkers and the posterior geometry. This method is well suited for multi-modal posteriors of moderate dimensions with unknown complex geometries and requires minimal tuning.

We contrast these Bayesian methods using several maximum likelihood estimation approaches. For these algorithms, the goal is to minimize the negative log-likelihood (i.e., maximize the log-likelihood) using numerical optimization strategies that generate a single estimate of the parameters that are the most likely. We perform randomized hyper-parameter tuning on each algorithm before minimizing the negative log-likelihood function: differential evolution[28], basin hopping[30], dual annealing [14], Nelder Mead [21], Powell [23], conjugate gradient [9], limited-memory Broyden–Fletcher Goldfarb–Shanno with box constraints (L-BFGS-B) [19], constrained optimization by linear approximations (COBYLA) [24], direct search [17] and sequential least squares programming (SLSQP) [18]. These methods, while not exhaustive, provide a survey of local and global optimizers [26, 15] covering a wide breadth of algorithmic strategies, and serve as useful baseline models to compare our pipelines' performance.

Our results suggest that preconditioned Monte Carlo and differential evolution have the best overall performance for the conditions studied. Preconditioned Monte Carlo costs approximately ten times as much as differential evolution but generates a much richer set of information containing parameter uncertainties and correlations.

#### 8.1 Log-Likelihood Comparisons

We first compare the estimated maximum log-likelihoods across the 12D and 16D models described in the manuscript - using multiple replicas for each maximum likelihood estimation algorithm with Bayesian min/max ranges shown.

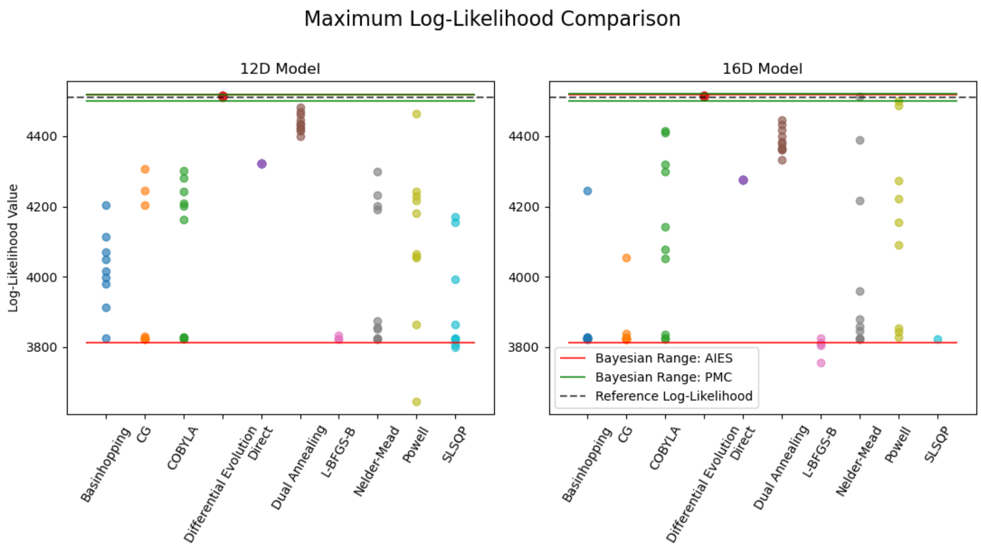


Figure 15: **Maximum Log-Likelihood Comparison - MLE.** The maximum log-likelihood values found for each MLE algorithm and replica. The reference log-likelihood is shown as a dashed line. Also, the Bayesian likelihood minimum and maximum values are shown for comparison. Most of the MLE algorithms fail to consistently estimate values near the reference value, or within the bounds of the PMC results. The exception is the differential evolution algorithm. We note the wide range of the AIES due to poor sampling convergence.

Next we examine the Bayesian log-likelihood distributions estimated from the PMC and AIES algorithms, with the best MLE results plotted for reference.

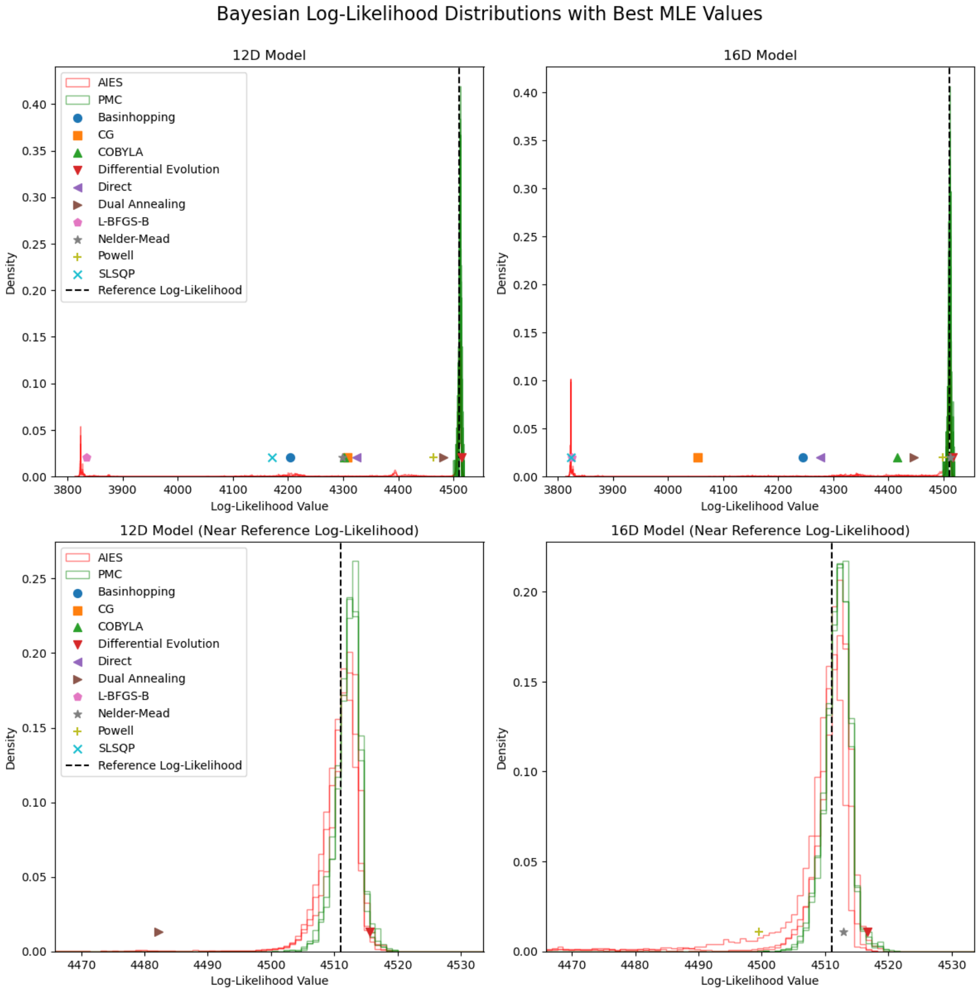


Figure 16: **Log-Likelihood Comparison - Bayes.** The log-likelihood distributions of each Bayesian algorithm and replica are shown, with the full range on the top row, and a reduced x-scale on the bottom row. The reference log-likelihood is shown as a dashed line, with markers denoting the best MLE estimates. The AIES algorithm appears to be stuck in local minimal as there are multiple modes at low log-likelihood values. Also, we find that the Bayesian inference methods find higher log-likelihood values than the MLE methods, with PMC sampling the maximum value. We note that that likelihoods larger than the reference value (based on ground-truth parameters) can occur due to noise in the data.

#### 8.2 Predicted Current Traces

We further assess these results using predictive analysis. Below we plot the predicted SSME-like current traces from all the ten MLE replicas and 100 randomly selected Bayesian posterior samples. We find that the differential evolution MLE method and PMC Bayesian method consistently predict datasets that fit the observed synthetic data well.

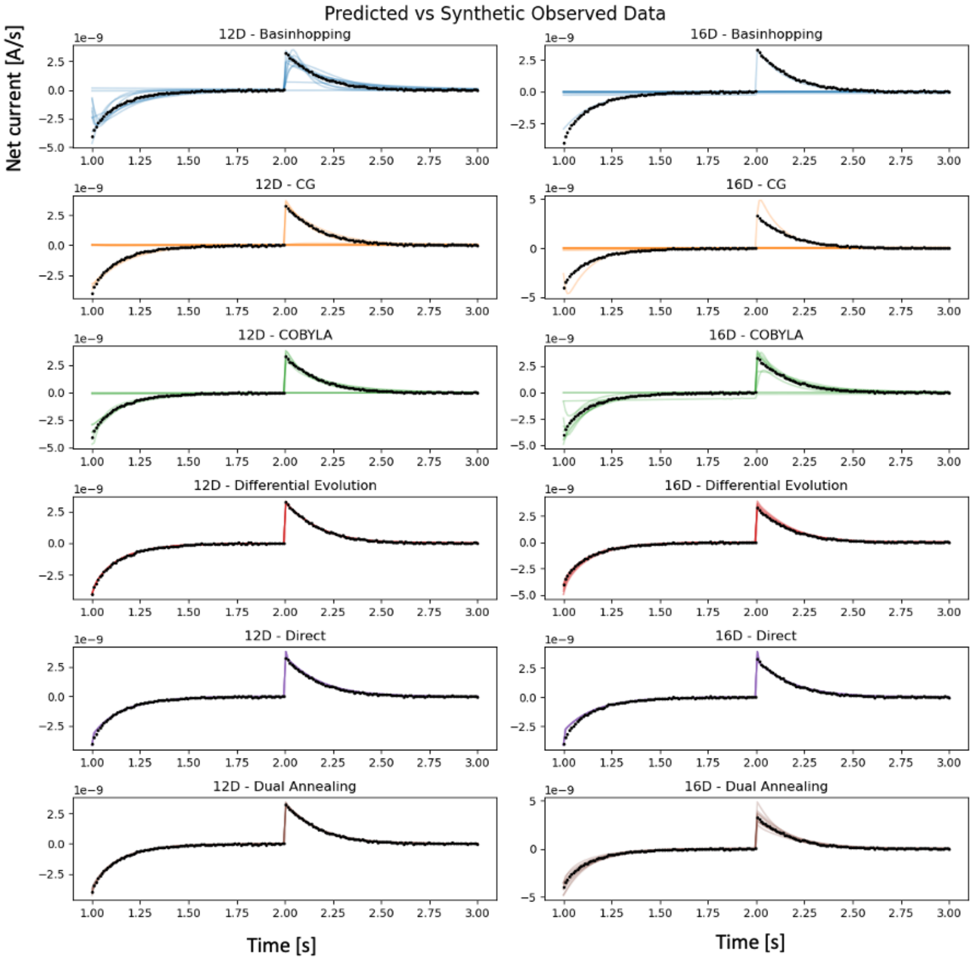


Figure 17: **Predicted Current Traces I.** The predicted current traces using the parameter sets from each MLE replica run plotted against the synthetic observed data (dots), for selected algorithms. Differential evolution and direct algorithms generate closely fit traces across both models.

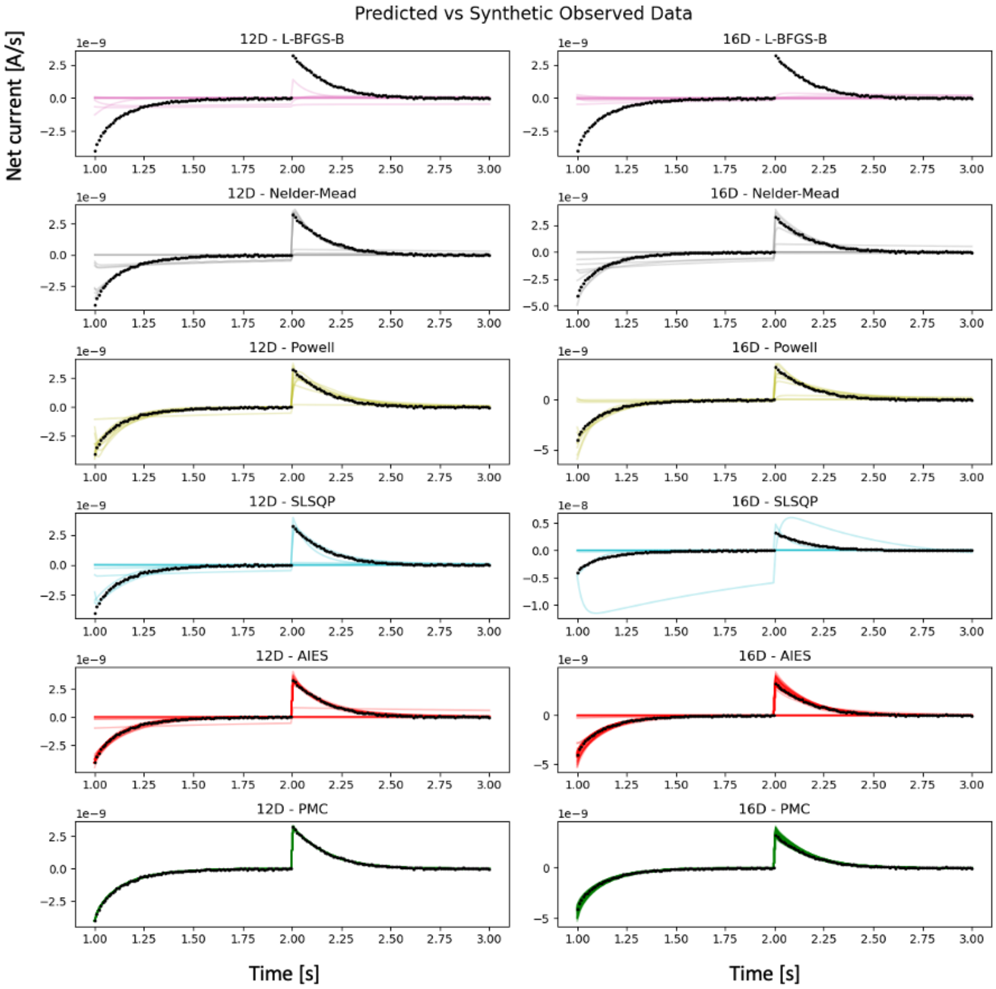


Figure 18: **Predicted Current Traces II.** The predicted current traces using the parameter sets from each MLE replica run plotted against the synthetic observed data (dots), for the remaining algorithms. For the Bayesian inference algorithms, 100 random parameter sets are chosen from the posterior and used to plot the current trace. Our pipeline using PMC generates closely fit parameter sets across both models.

#### 8.3 Computational Cost

In addition to evaluating the accuracy and consistency of each algorithm, we also examine the computational cost. We track the wall-clock run time in seconds (per replica), the number of log-likelihood evaluations used during the algorithm per replica, and the average number of log-likelihood evaluations per second (per replica). This is shown in Fig. 19. As expected the Bayesian inference methods incur the highest cost both in terms of run time and number of calculations used. The third most expensive algorithm was differential evolution, which used 1-2 orders of magnitude fewer log-likelihood calculations and wall clock time. However, we note that the Bayesian inference methods generate a full posterior that contains information for both parameter estimates as well as uncertainty quantification and parameter correlations. Furthermore, we find that the Bayesian inference methods have a comparable efficiency (log-likelihood calculations per second) as differential evolution.

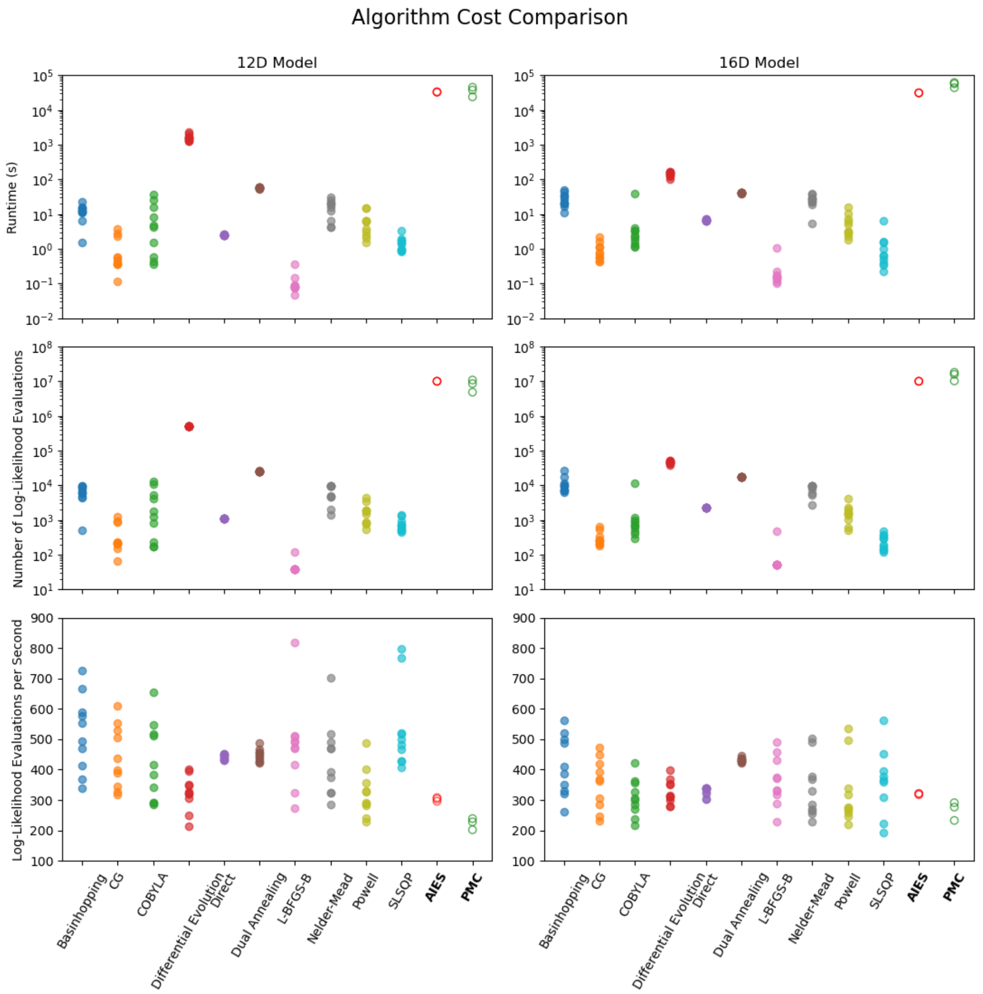


Figure 19: **Computational Cost Comparison.** The run time (top row), number of log-likelihood evaluations (middle row), and number of log-likelihood evaluations per second (bottom row) are shown for all the algorithms tested. The Bayesian inference methods AIES and PMC have the longest run time and number of log-likelihood evaluations, followed by differential evolution. We note that the Bayesian inference methods do not only estimate parameters, but also generate a full posterior distribution, and have a similar efficiency (log-likelihood calculations/sec) as differential evolution.

#### 8.4 Bayesian Marginal Posterior Analysis

Finally, we examine the posterior distributions for both the PMC and AIES methods, and both the 12D and 16D models. We find that a general agreement between the replicas for the PMC sampler, but not the AIES sampler, suggesting that AIES has not converged. The marginal posterior distributions are shown below:

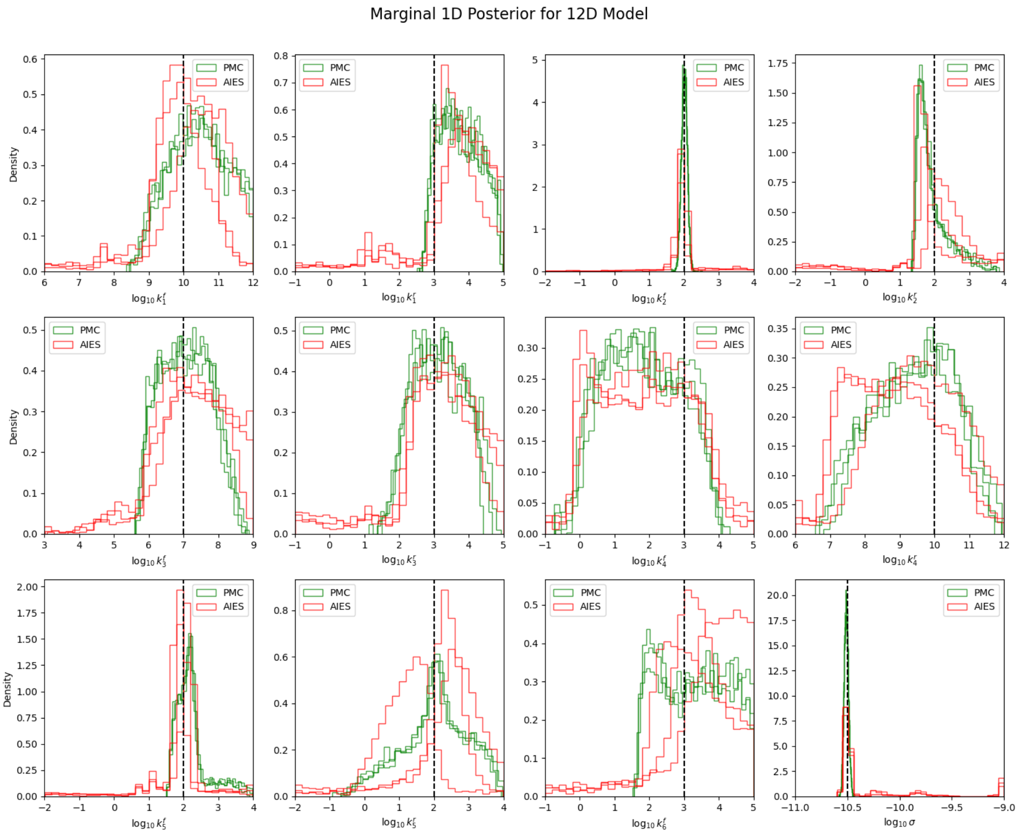


Figure 20: **Marginal Posteriors for 12D Model using PMC and AIES.** The posteriors from both methods cover the synthetic reference values (dashed lines). In contrast to PMC, the AIES replica distributions have increased variance between themselves which suggests that the sampler hasn't fully converged.

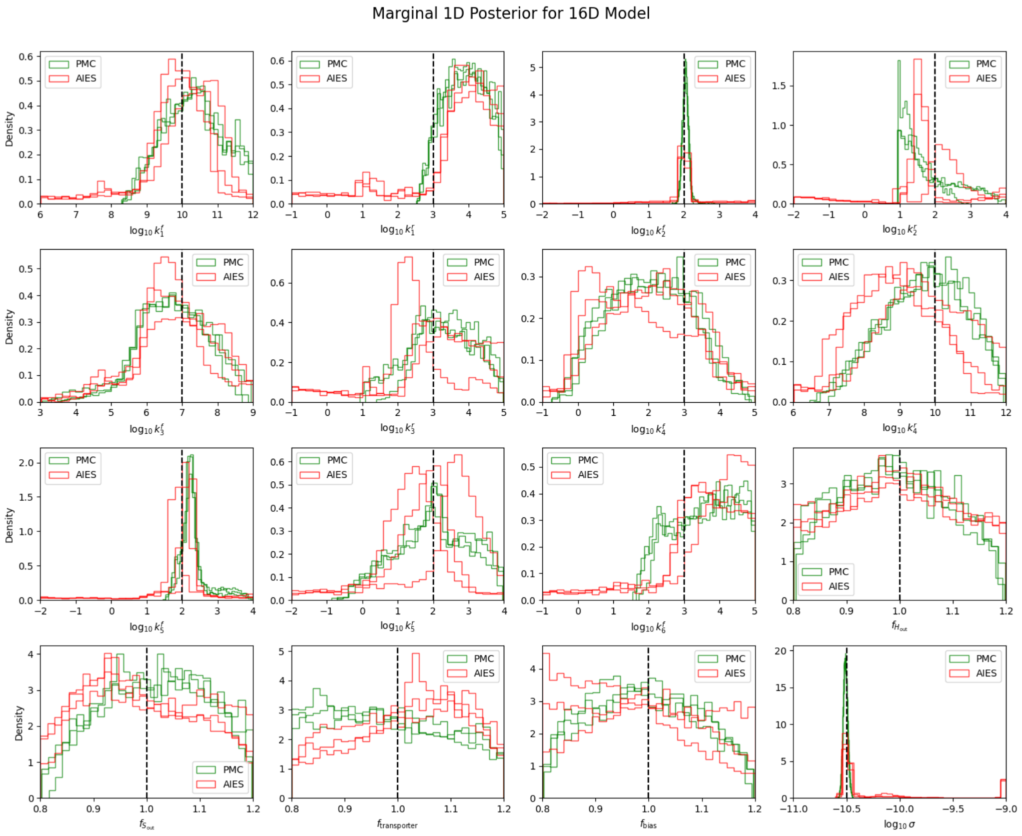


Figure 21: **Marginal Posteriors for 16D Model using PMC and AIES.** The posteriors from both methods cover the synthetic reference values (dashed lines). In contrast to PMC, the AIES replica distributions have increased variance between themselves which suggests that the sampler hasn't fully converged.
